# Remote immune processes revealed by immune-derived circulating cell-free DNA

**DOI:** 10.1101/2021.09.13.460029

**Authors:** Ilana Fox-Fisher, Sheina Piyanzin, Bracha-Lea Ochana, Agnes Klochendler, Judith Magenheim, Ayelet Peretz, Netanel Loyfer, Joshua Moss, Daniel Cohen, Yaron Drori, Nehemya Friedman, Michal Mandelboim, Marc E. Rothenberg, Julie M. Caldwell, Mark Rochman, Arash Jamshidi, Gordon Cann, David Lavi, Tommy Kaplan, Benjamin Glaser, Ruth Shemer, Yuval Dor

## Abstract

Blood cell counts often fail to report on immune processes occurring in remote tissues. Here we use immune cell type-specific methylation patterns in circulating cell-free DNA (cfDNA) for studying human immune cell dynamics. We characterized cfDNA released from specific immune cell types in healthy individuals (N=242), cross sectionally and longitudinally. Immune cfDNA levels had no individual steady state as opposed to blood cell counts, suggesting that cfDNA concentration reflects adjustment of cell survival to maintain homeostatic cell numbers. We also observed selective elevation of immune-derived cfDNA upon perturbations of immune homeostasis. Following influenza vaccination (N=92), B-cell-derived cfDNA levels increased prior to elevated B-cell counts and predicted efficacy of antibody production. Patients with Eosinophilic Esophagitis (N=21) and B-cell lymphoma (N=27) showed selective elevation of eosinophil and B-cell cfDNA respectively, which were undetectable by cell counts in blood. Immune-derived cfDNA provides a novel biomarker for monitoring immune responses to physiological and pathological processes that are not accessible using conventional methods.

## INTRODUCTION

Circulating biomarkers for monitoring inflammatory or immune responses are an essential part of diagnostic medicine and an important tool for studying physiologic and pathologic processes. These include, among others, counts of specific immune cell types in peripheral blood, RNA expression profiles in blood cells(Maas et al., 2002; Tuller et al., 2013), and levels of circulating proteins such as CRP(Gabay and Kushner, 1999; Sproston and Ashworth, 2018). A major limitation of circulating immune cell analysis is that it often fails to report on immune processes taking place in remote locations. Conversely, CRP and similar proteins do reflect the presence of tissue inflammation but are highly non-specific with regard to tissue location and the nature of inflammatory process(Gabay and Kushner, 1999).

Dying cells release nucleosome-size fragments of cell-free DNA (cfDNA), which travel transiently in blood before being cleared by the liver(Heitzer et al., 2019). Analysis of the sequence of such fragments is emerging as a powerful diagnostic modality. Liquid biopsies using cfDNA have been applied to reveal the presence of mutations in a fetus as reflected in maternal cfDNA(Bianchi et al., 2014; Christina Fan et al., 2012; Lo et al., 1997), identify and monitor tumor dynamics via the presence of somatic mutations in plasma(Wan et al., 2017), and detect the rejection of transplanted organs when the levels of donor-derived DNA markers are elevated in recipient plasma(De Vlaminck et al., 2015, 2014). More recently, we and others have shown that tissue-specific DNA methylation patterns can be used to determine the tissue origins of cfDNA, allowing to infer cell death dynamics in health and disease even when no genetic differences exist between the host and the tissue of interest(Cheng et al., 2019; Lehmann-Werman et al., 2016)

Although the majority of cfDNA in healthy individuals is known to originate in hematopoietic cells(Lehmann-Werman et al., 2016; Moss et al., 2018b; Sun et al., 2015), it has often been regarded as background noise, against which one may look for rare cfDNA molecules released from a solid tissue of interest. We hypothesized that identification of immune cell-derived cfDNA could open a window into immune and inflammatory cell dynamics, even in cases where peripheral blood counts are not informative. Here we describe the development of a panel of immune cell type-specific DNA methylation markers, and the use of this panel for cfDNA-based assessment of human immune cell turnover in health and disease. We show that immune cell cfDNA measurement can provide clinical biomarkers in multiple disease and treatment conditions, otherwise undetectable by cell subset enumeration in blood.

## RESULTS

### Identification of cell type-specific DNA methylation markers for immune cells

Using a reference methylome atlas of 32 primary human tissues and sorted cell types(Moss et al., 2018b), we searched for CpG sites that are hypomethylated in a specific immune-cell type and hypermethylated elsewhere. Consistent with the fundamental role of DNA methylation as a determinant of cell identity, we identified dozens of uniquely hypomethylated CpG sites for most cell types examined, qualifying these as biomarkers for DNA derived from a given cell type (**Figure 1A**). Based on this *in silico* comparative analysis, we selected for further work 17 different CpG sites, whose combined methylation status could distinguish the DNA of 7 major immune cell types: neutrophils, eosinophils, monocytes, B-cells, CD3 T-cells, CD8 cytotoxic T-cells, and regulatory T-cells (Tregs). For each marker CpG we designed PCR primers to amplify a fragment of up to 160bp flanking it, considering the typical nucleosome size of cfDNA molecules. Amplicons included additional adjacent CpG sites, to gain enhanced cell type specificity due to the regional nature of tissue-specific DNA methylation(Lehmann-Werman et al., 2016). We then established a multiplex PCR protocol, to co-amplify all 17 markers from bisulfite-treated DNA followed by next generation sequencing for assessment of methylation patterns(Neiman et al., 2020). Methylation patterns of amplified loci across 18 different human tissues validated the patterns inferred from *in silico* analysis and supported the ability of this marker cocktail to specifically identify the presence of DNA from each of the seven immune cell types (**Figure 1B**). We also assessed assay sensitivity and accuracy via spike-in experiments. We mixed human leukocyte DNA with DNA from the HEK-293 human embryonic kidney cell line and used the methylation cocktail to assess the fraction of each immune cell type. The markers quantitatively detected the presence of DNA from specific immune cell types even when blood DNA was diluted 10-20 fold (**Figure 1C and Supplementary Figure S1**). These findings establish specificity, sensitivity and accuracy of the methylation marker cocktail for detection of DNA derived from the 7 selected immune cell types.

**Figure 1:**
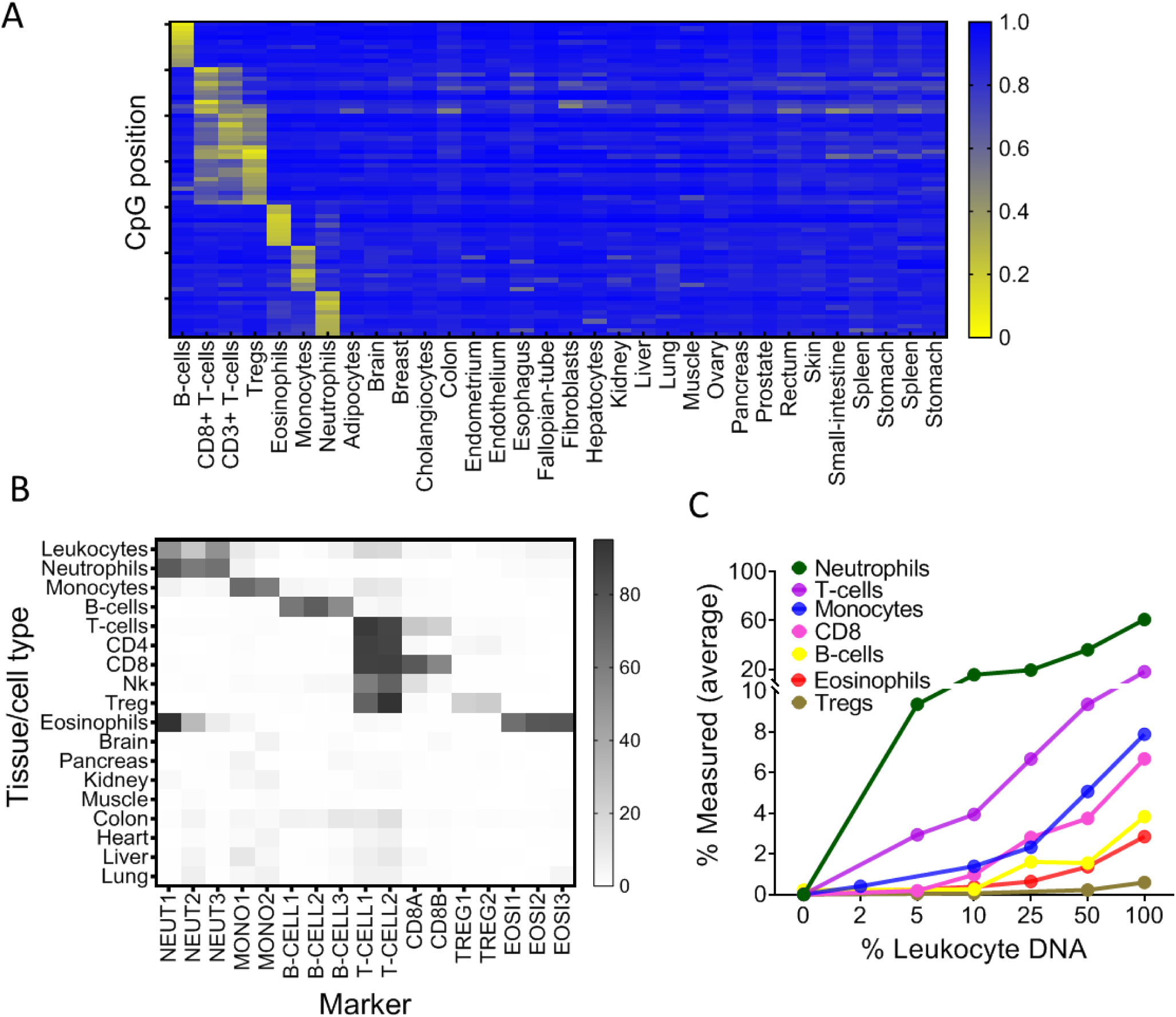
Identification of specific immune-cell DNA methylation markers. **A**, Methylation atlas, based on Illumina 450K arrays, composed of 32 tissues and sorted cells (columns). For each immune cell type we chose the top 10 CpGs that are hypomethylated (yellow) in the specific immune cell type and hypermethylated (blue) in other tissues and cells. This yielded 70 cell specific CpG sites (rows) for 7 different immune-cell subtypes– B-cells, CD8 cytotoxic T-cells, CD3 T-cells, regulatory T-cells, eosinophils, monocytes and neutrophils. **B,** methylation patterns of 17 loci, selected from the 70 shown in panel A, based on the presence of multiple adjacent hypomethylated CpGs within an amplicon of up to 160bp. Each methylation marker (columns) was assessed using genomic DNA from 18 different tissues and cell types (rows). All 17 markers were amplified in one multiplex PCR reaction. **C,** Spike-in experiments assessing assay sensitivity. Human leukocyte DNA was mixed with DNA from HEK-293 cells (human embryonic kidney cells) in the indicated proportions. Colored lines show the inferred proportion of DNA from each immune cell type using markers specific to neutrophils (NEUT1, NEUT2, NEUT3), monocytes (MONO1, MONO2), eosinophils (EOSI2, EOSI3), B-cells (B-CELL1, B-CELL2, B-CELL3), CD3-T-cells (T-CELL1, T-CELL2), CD8 cytotoxic T cells (CD8A, CD8B), and regulatory T cells (TREG1, TREG2).

### Immune cfDNA reflects cell turnover rather than counts of circulating blood cells

To validate assay accuracy, we applied the immune cell methylation markers to genomic DNA of blood cells, expecting to observe signals that agree with cell ratios as determined by Complete Blood Counts (CBC). We obtained 392 blood samples from 79 healthy individuals at different time points, and simultaneously tested for CBC, and methylation marker cocktail both in DNA extracted from whole blood as well as in cfDNA extracted from plasma. Theoretically, immune-cell cfDNA could be a mere reflection of the counts of each cell type (for example, if it is released mostly from blood cells that have died during blood draw or preparation of plasma). In such a case, immune cfDNA should correlate well with CBC (and with immune methylation markers measured in genomic DNA from whole blood). Alternatively, if cfDNA reflects cell death events that took place *in vivo*, the correlation to cell counts is expected to be weaker.

Comparing the CBC to DNA methylation pattern, we observed a strong correlation between assessments of specific cell fractions in the two methods (Pearson’s correlations; r=0.67-0.83, P-value<0.0001, **Figure 2A and Supplementary Figure S2**), supporting validity of the methylation assay for identifying fractions of DNA derived from specific immune cell types, consistent with previous findings(Baron et al., 2018). However, comparison of cfDNA methylation markers in plasma to blood DNA methylation markers and CBC, we observed that proportion of cfDNA from specific immune cell types did not correlate with the proportion of the same markers in circulating blood cells and with CBC (Pearson’s correlations; r=0.14-0.53, **Figure 2B-C and Supplementary Figure S2**). These findings suggest that immune cfDNA levels are the result of biological processes beyond immune cell counts.

**Figure 2:**
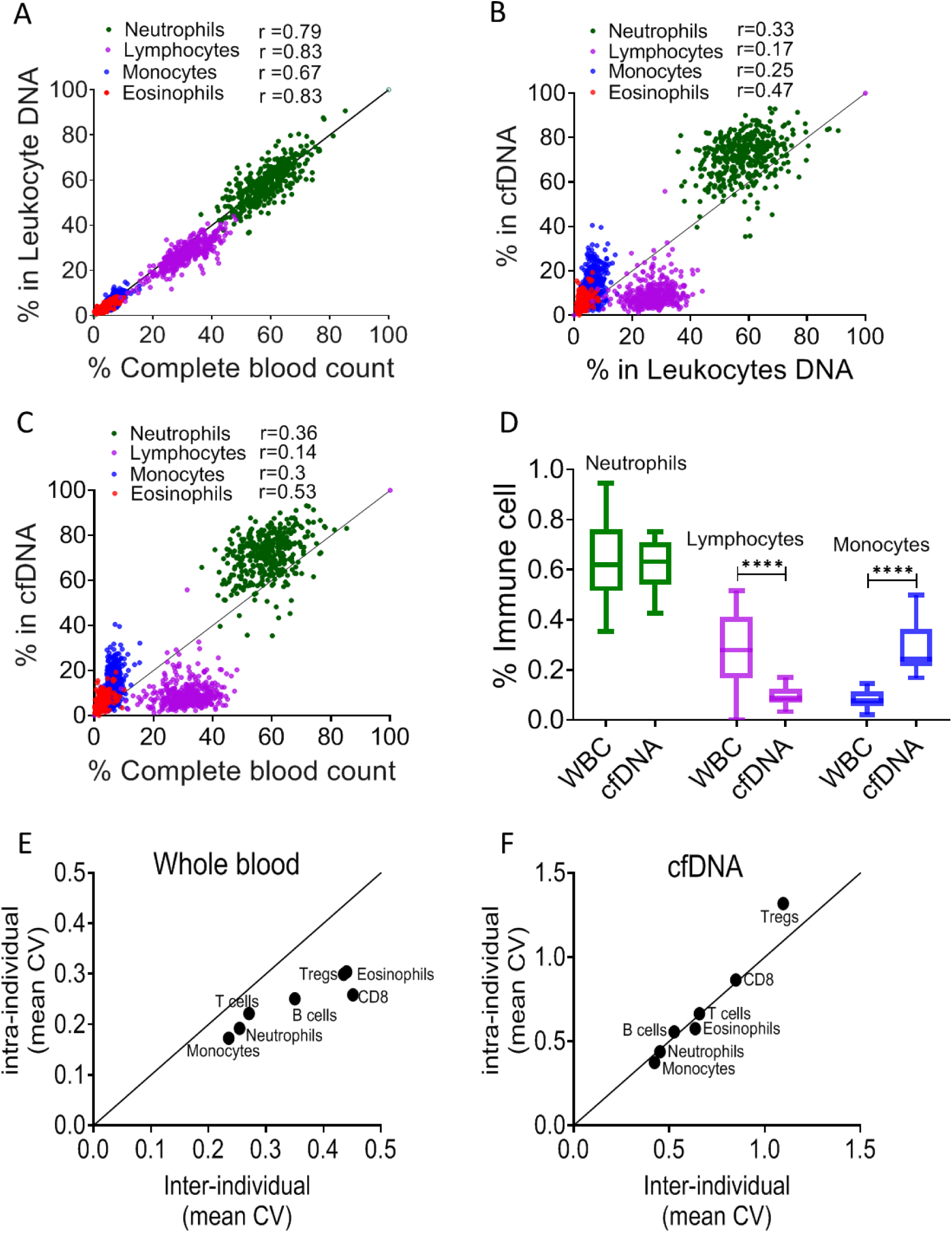
immune cfDNA methylation markers distribute differently than circulating immune cells and reflect immune cell turnover. **A**, levels of immune methylation markers in genomic DNA extracted from whole blood, versus Complete Blood Counts (CBC) from same donors. A total of 392 plasma and blood samples were obtained from 79 healthy donors (47 females, 32 males; age range 20-73). Shown are Pearson’s correlations; P-value<0.0001. **B,** cfDNA methylation versus whole blood methylation of the same donors. **C**, cfDNA methylation versus CBC of the same donors. Note that cfDNA proportions of immune cells differ from the proportions of these cell types in peripheral blood. **D,** deconvolution of cfDNA and blood cell (WBC) methylomes generated using whole genome bisulfite sequencing (WGBS) of 23 healthy donors. Note under-representation of lymphocyte DNA and over-representation of monocyte DNA in cfDNA compared with blood DNA (lymphocytes, p-value=0.0021; monocytes, p-value=0.0005, Kruskal-Wallis). **E-F,** XY scatter plots showing the average of inter-individual coefficient of variation (X-axis) and intra-individual coefficient of variation (Y-axis) for each immune cell type in whole blood (**E**) and in cfDNA (**F**) based on methylation markers. Black line represents perfect correlation between inter- and intra-individual dual variation. Dots below the black line reflect greater inter-individual variation and dots that are above reflect greater intra-individual variance. A smaller intra-individual variation in whole blood suggests a setpoint for proportions of blood cell types in each individual. By contrast, cfDNA levels of immune markers vary similarly within and between individuals.

We reasoned that over- or under-representation of DNA fragments from a specific immune cell type in plasma compared with blood counts most likely result from differences in cell turnover. The concentration of cfDNA from a given cell type should be a function of the total number of cells that have died per unit time (turnover rate), which is derived from the number of cells and their lifespan:

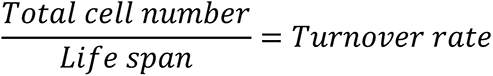

The larger the population of a given cell type (both circulating and tissue-resident), the more cfDNA it will release; similarly, the shorter is lifespan, the more DNA will be released to plasma. The cfDNA findings were consistent with this model. For example, the fraction of lymphocyte cfDNA in plasma was always smaller than the fraction of lymphocyte DNA in circulating blood cells or the fraction of lymphocytes in CBC (**Figure 2B-C and Supplementary Figure S2**), in agreement with the long half-life of lymphocytes compared with other blood cell types(Macallan et al., 2005; Michie et al., 1992). Conversely, the fraction of monocyte cfDNA was larger than the fraction of monocyte DNA in genomic DNA from whole blood or the fraction of monocytes in CBC (**Figure 2B-C**), consistent with the shorter half-life of monocytes(Patel et al., 2017).

To further examine the relative presence of immune cfDNA in plasma and whole blood, we employed an independent set of samples and an independent technology to measure and interpret methylation markers. Specifically, we performed deconvolution(Moss et al., 2018b) of methylomes obtained by whole genome bisulfite sequencing (30x coverage), on genomic DNA from whole blood, and matched plasma cfDNA from 23 healthy donors (see Methods). This analysis, based on genome-wide methylation patterns, also revealed that lymphocyte and monocyte cfDNA was under- and over-represented, respectively, relative to the abundance of DNA from these cells in blood (lymphocytes, p-value=0.002; monocytes, p-value=0.0005, Kruskal-Wallis) (**Figure 2D**).

These findings support the exciting idea that cfDNA levels from a given immune cell type integrate total cell number and the lifespan of that cell type, and can provide information on processes not evident from circulating cell counts. For example, if the level of cfDNA from a specific immune cell type increases while the circulating counts of this cell type are unchanged, this can indicate either growth in the size of a tissue-resident population, or increased cell turnover, both of which are important immune parameters that cannot be easily obtained otherwise. Below we provide evidence that such information can be extracted following perturbations of immune homeostasis.

### Estimation of immune cfDNA baseline in healthy individuals

To define the baseline levels of immune-derived cfDNA in healthy individuals, we collected and tested plasma samples from 227 healthy donors (males and females, ages 1-85 years). Consistent with our previous plasma methylome analysis(Moss et al., 2018a) , we found that neutrophils were the main source of immune-derived cfDNA (Mean=390, range 10-1064 genome equivalents [GE] /ml), followed by monocytes (Mean=101, range 6-233 GE/ml), eosinophils (Mean=38,range 0-111 GE /ml) and lymphocytes (T cells, Mean=30, range1-79; B cells, Mean=17,range 3-42; CD8 T-cells, Mean=8,range 0-27; Tregs, Mean=2, range 0-9 GE/ml) (**Supplemental Figure S3)**. These data chart the normal range of cfDNA concentrations from specific immune cell types, including age and gender characteristics, against which pathologic deviations can be identified (see below).

We also conducted a longitudinal study, to understand how immune-derived cfDNA is changing over time in the same individual. We collected weekly blood samples from 15 healthy donors over a period of 6 weeks. For each sample we obtained CBC and measured immune DNA methylation markers in genomic DNA of blood cells and in plasma cfDNA. We then calculated the coefficient of variation (CV) among CBC, blood methylation markers and cfDNA methylation markers, within and between individuals. In circulating blood cells, the inter-individual variation in immune methylation markers and CBC was always higher than the intra-individual variation in these markers (**Figure 2E and Supplementary Figure S3**). This is consistent with previous reports that blood cell counts among individuals are more similar to themselves than to others, indicating distinct set-points per person for the total number of specific immune cell types circulating in blood(Alpert et al., 2019; Carr et al., 2016). Strikingly, cfDNA values of the same immune methylation markers varied to the same extent among samples of the same individual and among samples of different individuals (**Figure 2F**). This argues that unlike cell counts, cfDNA of immune cells has no individual set-point. Rather, cfDNA levels appear to reflect homeostatic maintenance of cell number, whereby cell birth and death are modulated to maintain a desired cell count.

### Elevation of B-cell derived cfDNA after influenza vaccination precedes changes in cell counts and correlates with specific antibody production

We hypothesized that upon perturbations of the immune system, cfDNA markers will reveal information about immune cell dynamics that is not present in peripheral blood cell counts, for example extensive cell death during the process of affinity maturation, which repeatedly selects for B cell clones with increased antibody-target affinity. To test this hypothesis, we examined longitudinal blood samples from healthy individuals who received an annual quadrivalent influenza vaccination(Nakaya et al., 2011; Voigt et al., 2018). The influenza vaccine response is mediated mostly by the humoral immune system (B cells) aided by CD4 T-cells(Gage et al., 2018). Changes in circulating cell counts occur a week after vaccination, reflecting processes such as plasma cell formation (Victora and Wilson, 2015). cfDNA responses to vaccination were not previously reported. We recruited 92 healthy volunteers (age range 20-73, mean age 37.4) who received the vaccination in 2018 or 2019. From each volunteer we obtained blood samples a day before vaccination (day zero, D0), and at day 3, 7 and 28 post-vaccination. Consistent with previous reports, B cell counts (measured by methylation analysis of DNA from whole blood) were moderately but significantly elevated on day 7, and persisted to day 28 (p-value=0.0048, Kruskal-Wallis) (**Figure 3A**)(Li et al., 2012). Surprisingly, B-cell derived cfDNA levels increased as early as day 3, peaked on day 7 and returned to baseline levels on day 28 (p-value<0.0001, Kruskal-Wallis) (**Figure 3B,D**), suggesting that cfDNA reveals an early increase in the turnover of B-cells following vaccination, which is not portrayed in circulating B-cells. We observed a similar trend in the ratio of B-cell cfDNA to B-cell counts in each individual (**Figure 3C**, p-value=0.016, Kruskal-Wallis, B-cell counts calculated from methylation markers in whole blood). Of note, this response was specific to B-cell-derived cfDNA; total cfDNA levels did not change over the time course of vaccination, nor did cfDNA levels of other immune cell types (**Supplementary Figure 4S)**. Taken together, this strengthens evidence that cfDNA changes reflect processes beyond alterations in absolute circulating cell counts in a cell-specific manner.

**Figure 3:**
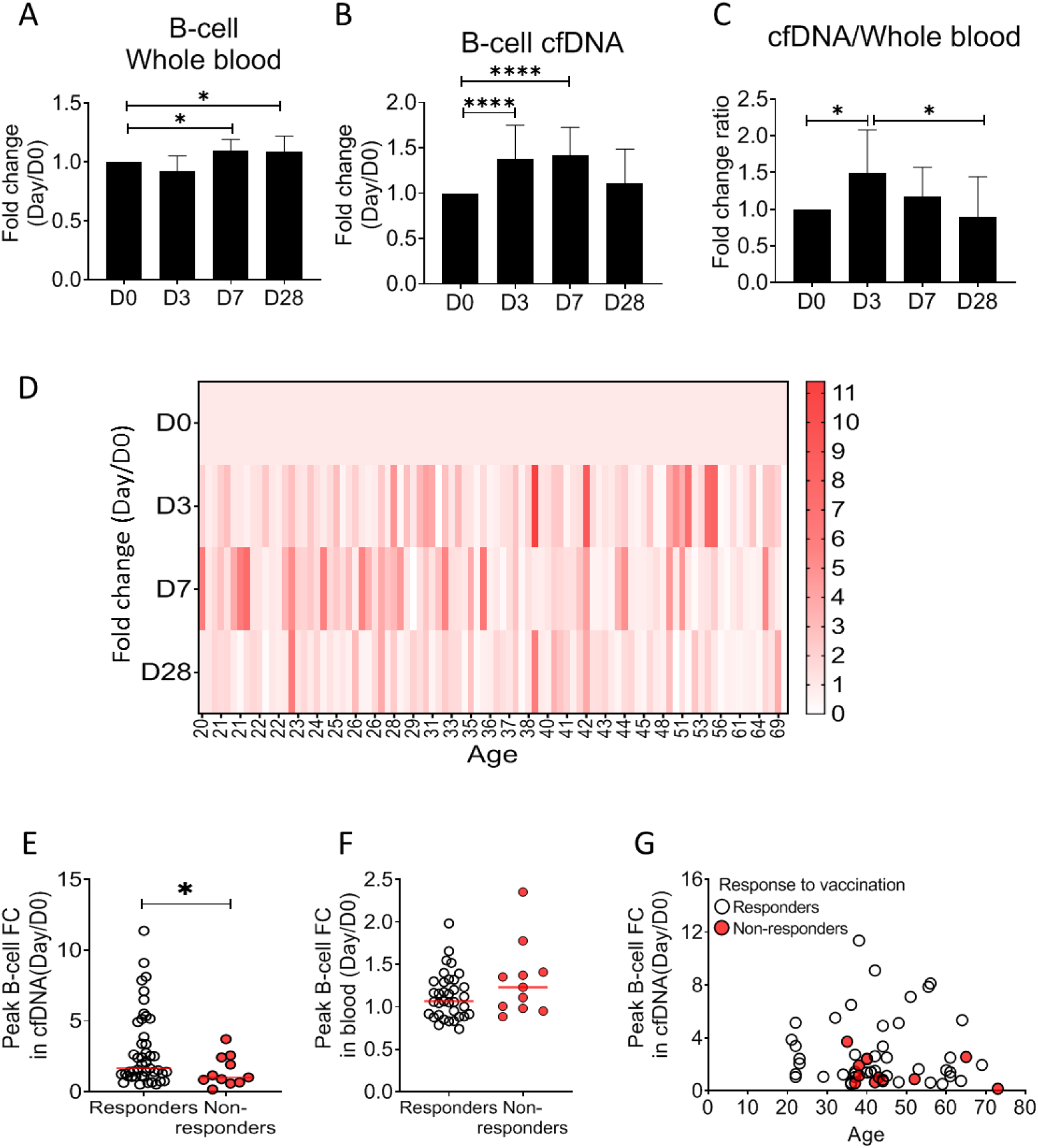
Elevation of B-cell derived cfDNA after influenza vaccination precedes changes in cell counts and reflects efficacy of response to vaccination. Plasma, serum, and blood samples were obtained from 92 healthy donors receiving the influenza vaccination (55 females, 37 males, age range 20-73y). **A,** circulating B-cells, assessed using methylation markers, are elevated on day 7 (p-value=0.03) and 28 (p-value=0.021) compared to baseline (Kruskal-Wallis). **B,** B-cell derived cfDNA markers where normalized to the levels of each individual at baseline (D0) and represented as fold change. B-cell derived cfDNA is elevated compared to baseline on day 3 and day 7 after influenza vaccination (p-value<0.0001, Kruskal-Wallis). **C,** the ratio of B-cell derived cfDNA and B-cells in blood was calculated for each individual. On day 3 the ratio was significantly higher than at baseline (p-value=0.016). Bars indicate the median and error bars indicate the confidence interval. **D**, a heat map showing the change of B-cell derived cfDNA in each individual following vaccination relative to baseline. Donors are ordered by their age. **E,** serum antibody titers were used to divide donors into responders (n=45) who have developed antibodies following vaccination, and non-responders (n=11) who have not. Graph shows the maximal elevation (fold change [FC] from baseline) in B-cell derived cfDNA that was recorded for each donor. (P-value=0.045, Mann-Whitney). **F,** Maximal elevation in B-cell counts in blood based on methylation markers does not differ between responders and non-responders. P-value=0.2, Mann-Whitney test. **G,** XY Scatter plot for peak B-cell cfDNA elevation as a function of age. Non- responders (colored with red) tend to be older and show reduced elevation of B-cell cfDNA.

To ask if the elevation of B-cell cfDNA has functional significance in the development of an immune response, we obtained information on the production of antibodies. We classified all volunteers into responders or non-responders according to the hemagglutinin antibody titer measured at 28 days post-vaccination, and asked if B-cell cfDNA or B-cell counts correlated with antibody production. Strikingly, responders had a higher peak elevation of B-cell cfDNA relative to their pre-vaccination baseline levels compared with non-responders (p-value=0.044, Mann-Whitney) (**Figure 3E and** Supplementary **Figure S4**). Peripheral B-cell counts were not different between responders and non-responders (p-value=0.2, Mann-Whitney) (**Figure 3F**). It is well established that influenza vaccination is more effective in younger individuals(Del Giudice et al., 2015; Ranjeva et al., 2019; Siegrist and Aspinall, 2009; Wagner et al., 2018). To examine the relationship between age, antibody production and cfDNA we plotted the fold elevation of B-cell cfDNA from baseline as a function of donor age, and marked responders and non-responders Non-responders to vaccination in our cohort were all above 35 years and tended to have a minimal elevation of B-cell cfDNA above baseline even when compared to people in their age group (**Figure 3G and Supplementary Figure S4,** peak elevation of B cell cfDNA in responders vs non-responders p value=0.089), suggesting that B-cell cfDNA dynamics report on a biological process independent of age. We conclude that B-cell turnover (as reflected in B-cell cfDNA but not in B-cell counts) captures an early response of the immune system to influenza vaccination that predicts a functional outcome, suggesting cell-specific cfDNA could serve as a sensitive biomarker of functional immune changes.

### Selective elevation of eosinophil-derived cfDNA in patients with Eosinophilic Esophagitis

To test the hypothesis that immune-derived cfDNA can reveal pathologic inflammatory processes in remote locations, we studied patients with Eosinophilic Esophagitis (EoE). EoE is a chronic inflammatory disease characterized clinically by esophageal dysfunction, and histologically by eosinophil-predominant inflammation of the esophagus(Liacouras et al., 2011). Diagnosis of EoE requires an invasive endoscopic biopsy. Notably, most patients do not show peripheral eosinophilia(Aceves et al., 2007; Dellon et al., 2009). We analyzed blindly immune cfDNA markers in plasma samples from patients with active EoE (N=21), patients with EoE in remission (N=24) and healthy controls (N=14). Patients with active EoE had elevated levels of Eosinophil cfDNA (Mean=115 GE/ml) compared with both healthy controls (Mean=34 GE/ml, p-value =0.0056) and patients with inactive EoE (Mean=36 GE/ml, p-value=0.0003, Kruskal-Wallis), while other immune cfDNA markers were not elevated in active EoE patients (**Figure 4A,B and Supplemental Figure 5S**). The fraction of eosinophils in blood was not significantly elevated in EoE patients (**Figure 4C**, p-value=0.1, Kruskal-Wallis), consistent with restriction of eosinophil abundance to the esophagus and further supporting the idea that immune cfDNA is not a reflection of circulating immune cells.

**Figure 4:**
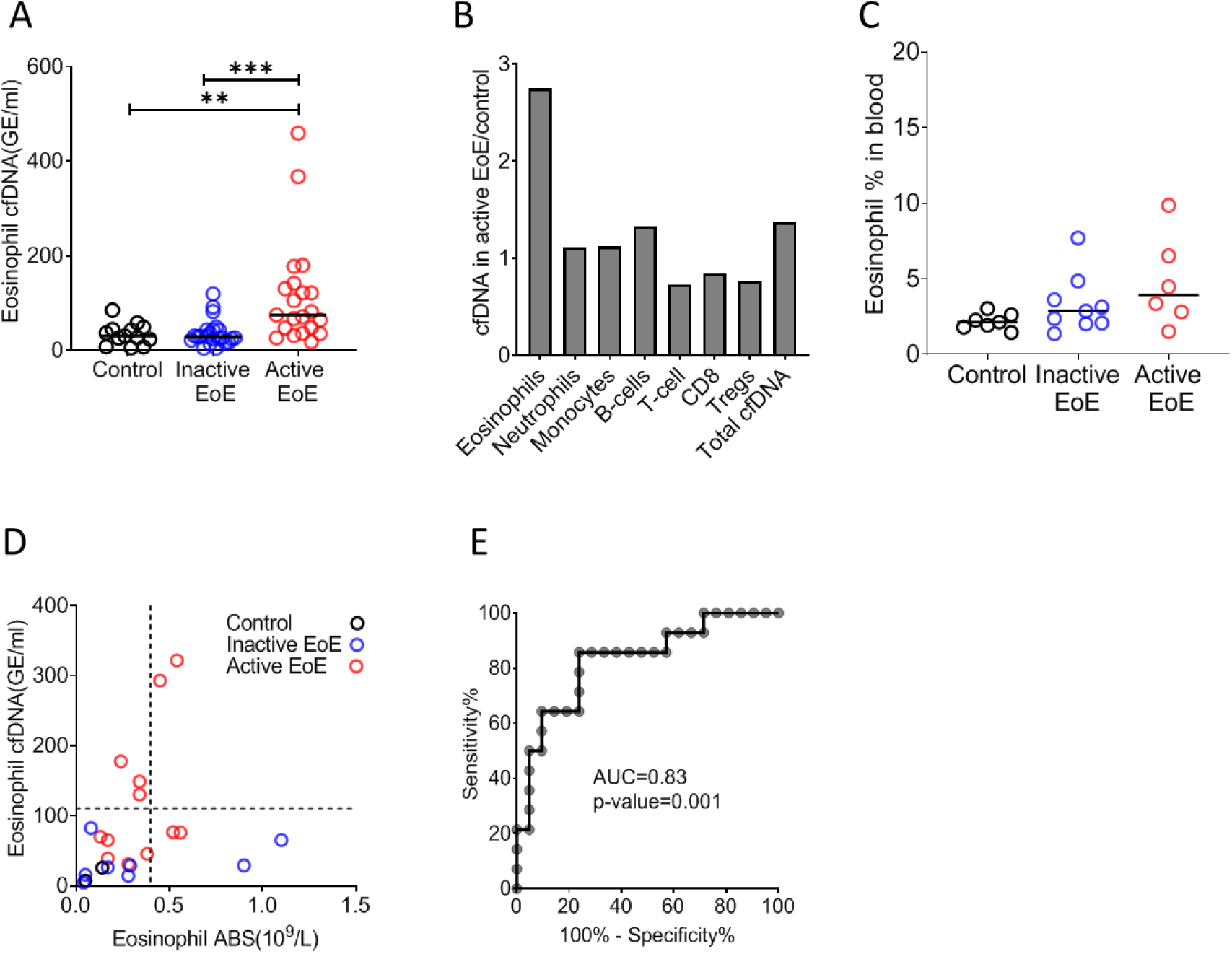
Selective elevation of eosinophil-derived cfDNA in patients with Eosinophilic Esophagitis. **A**, eosinophil derived cfDNA in Active EoE patients (n=21) compared with healthy controls (n=14, p-value=0.0056) and patients with EoE in remission (inactive EoE, n=24, pvalue= 0.0003, Kruskal-Wallis). **B**, differences between immune cfDNA populations in Active EoE patients and healthy controls (mean active EoE/ mean control). **C**, eosinophil DNA markers in whole blood of patients and controls (p-value=0.1, Kruskal-Wallis test). **D**, XY Scatter plot for eosinophil derived cfDNA levels vs. eosinophil absolute count (ABS) in blood. Dashed lines indicate healthy maximal baseline levels of eosinophil absolute counts in blood, and eosinophil-derived cfDNA in plasma. **E**, receiver operating characteristic (ROC) curve for the diagnosis of active EoE, using eosinophil cfDNA levels in plasma of healthy controls and patients with Active EoE.

Among a small subset of donors for which we had access to plasma, PBMC and CBC (12 active EoE, 8 inactive EoE, 3 controls), elevated eosinophil counts and elevated eosinophil cfDNA levels were observed in non-overlapping groups of EoE patients (elevated eosinophil counts in 4/12 patients with active EoE, 2/8 patients with inactive EoE, as previously reported(Dellon et al., 2009); elevated eosinophil cfDNA in 5/12 patients with active EoE), suggesting that counts and cfDNA reflect different biological processes (**Figure 4D**). Finally, we tested our ability to distinguish active EoE patients from healthy individuals, noting high specificity and sensitivity (**Figure 4E**, AUC 0.83, p-value = 0.001). These findings suggest that cell type-specific cfDNA can be used to detect clinical inflammation occurring in solid tissues that is not reflected in peripheral cell counts.

### B-cell derived cfDNA elevation in patients with B-cell lymphoma

Hematologic malignancies occurring in remote immune organs such as the bone marrow, spleen and lymph nodes are often undetectable in peripheral blood(Conlan et al., 1991) . We reasoned that increased turnover of cancer cells in such malignancies would release cfDNA molecules carrying methylation marks of the normal cell type from which the tumor originated, informing on tumor presence and dynamics. In addition, cell type-specific cfDNA markers could reveal collateral damage incurred by the tumor to normal adjacent cells(Ménétrier-Caux et al., 2019; Ray-Coquard et al., 2009). To test this idea we examined blood samples from patients with B-cell lymphoma, a disease which often requires imaging and invasive biopsies for diagnosis and monitoring(Barrington et al., 2014; Laurent et al., 2017). We studied plasma and blood cells from 17 newly diagnosed (treatment-naïve) B-cell lymphoma patients (diffuse large B-cell lymphoma, n=6; Hodgkin’s lymphoma, n=5; Follicular lymphoma, n=6) and age-matched healthy controls (**Data file S1**). Lymphoma patients (Mean=264.4 GE/ml) had dramatically elevated levels of B-cell derived cfDNA compared with controls (Mean=18.3 GE/ml, p-value<0.0001), while B cell counts in peripheral blood were actually decreased (control; Mean=0.162, Lymphoma; Mean=0.079, 10^9^/L, p-value=0.0059, Mann-Whitney) (**Figure 5A-C**). We observed that the level of B-cell cfDNA accurately distinguished B-cell lymphoma patients from healthy controls, much better than did B-cell counts (cfDNA, AUC=0.98, p-value<0.0001; B-cell counts, AUC=0.75, p-value=0.006; **Figure 5D,E)**. Total levels of cfDNA as well as the levels of other immune cfDNA markers were also elevated in lymphoma patients, consistent with reports on alterations in non-B-cells in lymphoma(Simone, 2013). We observed the strongest response in the levels of B-cell cfDNA (14.4 fold increase compared with controls), CD8 cytotoxic T-cells (10.7 fold) and Tregs (13.8 fold) (**Figure 5F, Supplemental Figure 6S**). Lymphocyte counts were decreased, such that the ratio of cfDNA to cell count for each cell type was dramatically elevated in lymphoma patients (**Figure 5G**). We did not observe a correlation between lymphoma type and immune cfDNA patterns, perhaps because of the small sample size. To validate these findings, we performed the analysis on plasma samples from a second, independent cohort of lymphoma patients and healthy controls. As in the first cohort, we observed elevated B cell cfDNA in patients (lymphoma n=10, Mean=1473 GE/ml; Control n=34, Mean=15 GE/ml, p-value<0.0001) (**Figure 5H**), accompanied by lower B cell counts and elevated T cell cfDNA (**Supplemental Figure S7)**.

**Figure 5:**
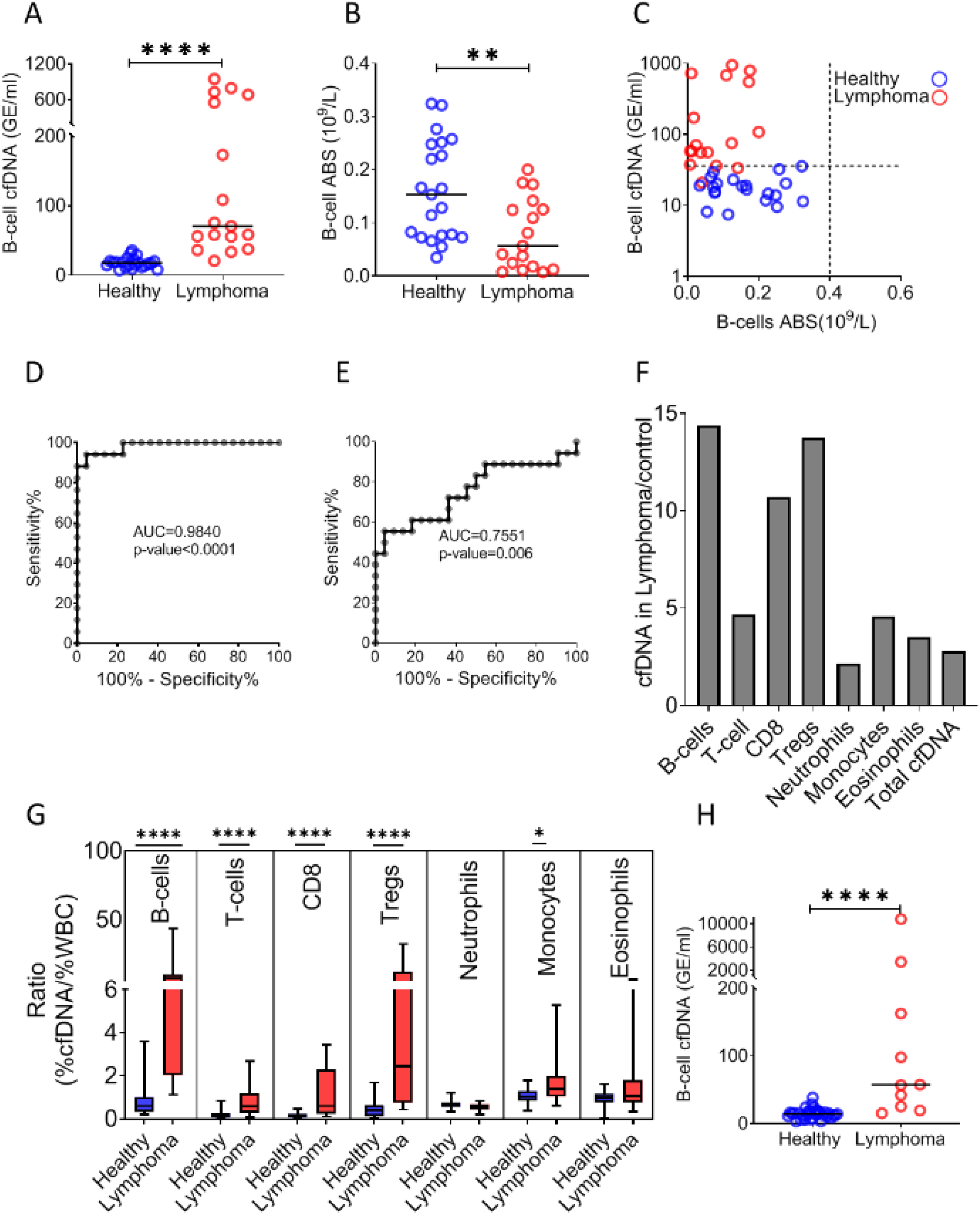
B-cell derived cfDNA elevation in patients with B-cell lymphoma. **A**, B-cell derived cfDNA in patients with lymphoma (n=17) compared with age-matched healthy controls (n=23, pvalue< 0.0001, Mann-Whitney). **B**, B-cell absolute counts in patients with lymphoma compared with age-matched healthy controls (p-value=0.0059, Mann-Whitney). **C**, XY Scatter plot for B-cell derived cfDNA levels versus B-cell absolute counts in blood. Dashed lines indicate healthy baseline levels of B-cell absolute counts in blood and B-cell cfDNA. **D**, ROC curve for the diagnosis of lymphoma based on B-cell cfDNA levels in healthy subjects and patients with B cell lymphoma. **E**, ROC curve for diagnosis of lymphoma based on B-cell counts. **F**, levels of immune cell type-specific cfDNA in lymphoma patients and healthy controls (mean lymphoma/ mean control). **G**, the ratio between the percentage of cfDNA from a given immune cell type and the percentage of cells from this population in blood according to CBC, in each donor among the healthy volunteers (n=23, blue bars) and patients with lymphoma (n=17, red bars). Boxes represent 25th and 75th percentiles around the median, whiskers span min to max. **H**, B-cell derived cfDNA in an independent cohort of 44 donors including 10 patients with lymphoma and 34 healthy controls (p-value<0.0001, Mann-Whitney).

These findings indicate that lymphoma growth causes an elevation in the levels of B-cell cfDNA. In addition, a massive loss of normal T cells leads to extensive release of cfDNA, potentially reflecting an immune response against the tumor or collateral damage. Taken together across all three conditions (influenza vaccination, EoE and lymphoma), immune cell dynamics in remote locations that are not evident in peripheral blood are detectable via cell-specific methylation markers in plasma.

## DISCUSSION

We describe a novel method for monitoring turnover dynamics of the human immune system, using cell type-specific cfDNA methylation markers. The assay opens a window into aspects of human immune and inflammation biology that are not reflected in blood cell counts or gene expression patterns. Specifically, the concentration of cfDNA derived from a given immune cell type is a function of the total number of cells of that type (circulating and remote pools, combined), the lifespan of this population, determinants of cfDNA release (e.g. efficiency of phagocytosis) and determinants of cfDNA clearance from plasma (e.g. liver uptake). While many of these parameters are typically unknown, in some cases cfDNA dynamics may allow to infer a change in cell turnover or in total cell number outside systemic circulation.

Since the method relies on highly stable methylation marks of cell identity(Dor and Cedar, 2018), it is expected to be universal, with the same markers allowing to accurately monitor immune cell dynamics in all individuals. While our current assay uses a panel of 17 methylation markers specific to 7 key immune cell types, future improvements (such as adding markers specific to other cell types) should increase the resolution of analysis to target essentially all immune cell types. We note that dynamic cellular states may involve changes in gene expression that do not involve reprogramming of DNA methylation patterns, representing a limitation of the approach.

Our cross-sectional and longitudinal analysis of immune cfDNA dynamics in healthy individuals begins to define the normal range among the population, an essential step towards using the assay for identifying with confidence deviations from health. More extensive characterization of immune cell cfDNA in healthy individuals is necessary to interpret trends that were revealed by our healthy cohorts. For example, we noticed lower levels of neutrophil cfDNA in adult females compared with adult males, suggesting that neutrophils in females live longer; we speculate that such differences in lifespan explain why women have a higher steady state neutrophil count(Bain and England, 1975) (and data not shown). Additional observations of healthy immune cfDNA dynamics that merit further investigation regard age-related changes, for example elevated monocyte cfDNA and reduced lymphocyte cfDNA in individuals older than 60. Finally, the intra- and inter-individual variation in immune cfDNA levels show that unlike blood cell counts, cfDNA levels vary wildly, apparently with no regulatory mechanism that attracts them to a certain setpoint typical to an individual. We propose that varying cfDNA levels reflect the action of regulated cell death as a mechanism by which the healthy body dynamically maintains homeostatic cell numbers within a desired range (model, **Figure 6**).

**Figure 6:**
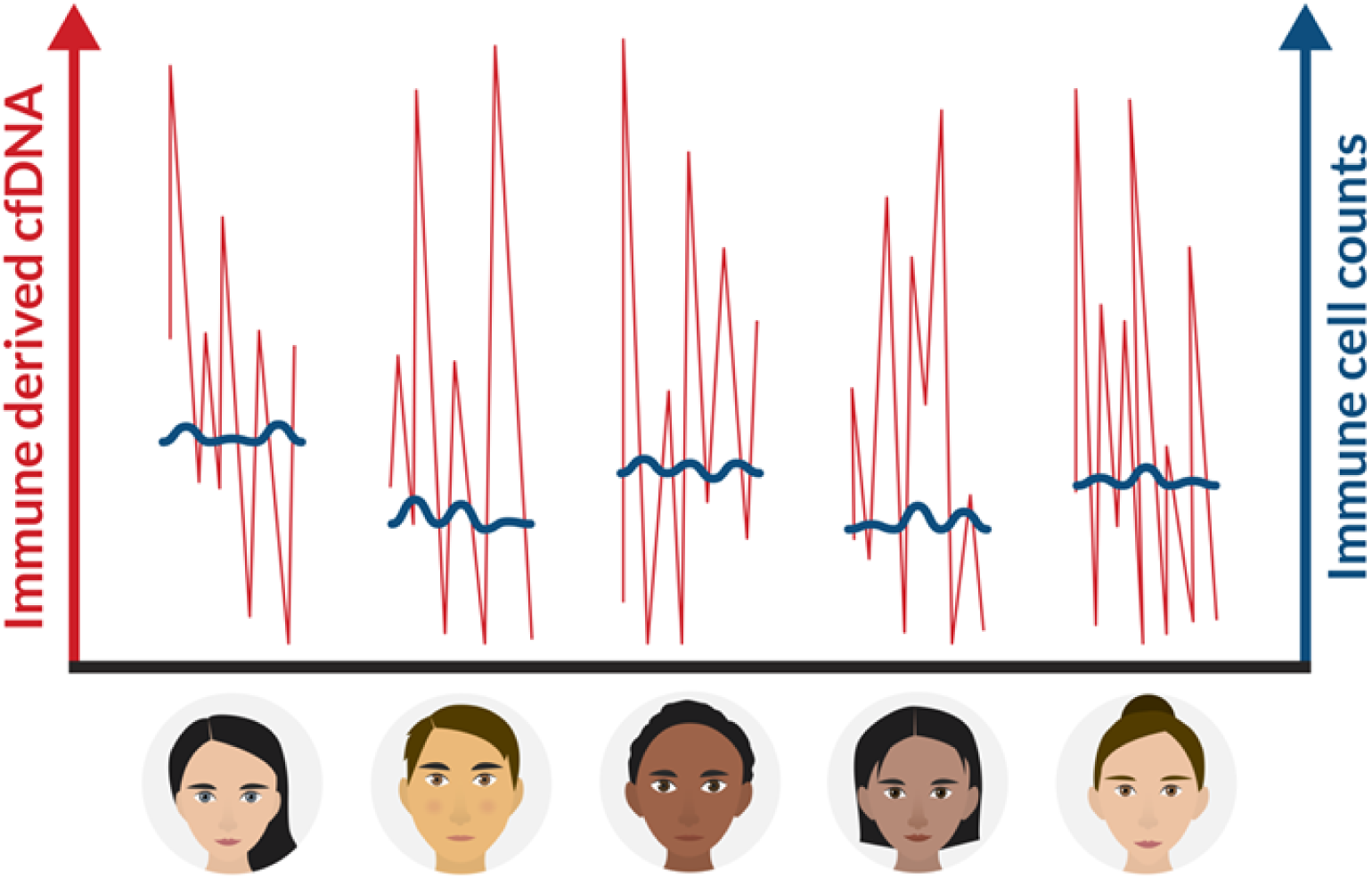
A schematic view of immune marker variance within individuals. Intra-individual variance of immune-cell count (blue) and immune-derived cfDNA (red) in multiple time points. Our findings suggest that while immune cell counts are stable and typical within an individual, immune cell cfDNA levels vary greatly, reflecting changes in cell turnover that help maintain the cell count set point.

Beyond the healthy baseline, we studied immune cfDNA dynamics in three settings of immune system perturbation. First, post influenza vaccination we identified an early elevation of B cell cfDNA, preceding an increase in circulating B-cell counts and showing a striking correlation to the effectiveness of antibody production, which was independent of the known age-related risk of non-responsiveness. We propose that elevated B-cell cfDNA reflects early stages in the successful response of B-cells to the vaccine, including the process of affinity maturation whereby large numbers of B-cells are generated and eliminated within lymph nodes as a result of insufficient binding to the target epitope. More work is needed to accurately define the population of B-cells that release cfDNA after vaccination, and to understand the physiological driver of this response. Practical applications may include an early indication for the success of experimental vaccination. Second, we examined immune cfDNA dynamics in EoE, a model for an inflammatory disease in which one tissue is damaged by infiltration of a specific immune cell population, while leaving a minimal mark on peripheral cell counts. cfDNA analysis revealed the selective elevation of eosinophil turnover in EoE, in some cases even when circulating eosinophil cell counts are unchanged. Larger scale studies are warranted to determine if eosinophil cfDNA can be a sufficiently sensitive and specific biomarker for assisting the clinical diagnosis and monitoring of EoE, ultimately relieving the need for invasive biopsies of the esophagus. Lastly, cfDNA dynamics in patients with B-cell lymphoma revealed the impact of disease on the turnover of B-cells and other immune cell types. As with EoE, cfDNA in lymphoma provides a systemic biomarker of immune processes taking place in remote locations. However in lymphoma, these processes include both tumor dynamics and host responses – either bystander effects (collateral damage) or an immune response to the tumor. Potential uses of immune methylation markers in this field include early diagnosis of hematologic malignancies, detection of minimal residual disease, and monitoring response to treatment. Beyond hematologic malignancies, immune-derived cfDNA dynamics can inform on the response to immune checkpoint inhibitors.

In summary, analysis of specific immune cell methylation markers in cfDNA allows for monitoring of human immune cell dynamics, providing temporal and spatial information not accessible via circulating cell counts. We propose that this novel tool can illuminate healthy and pathologic immune processes, including non-immune diseases having an inflammatory component such as cancer, rejection of transplanted organs, metabolic and neurodegenerative disease.

## MATERIALS AND METHODS

### Subject enrollment

This study was conducted according to protocols approved by the Institutional Review Board at each study site, with procedures performed in accordance with the Declaration of Helsinki. Blood and tissue samples were obtained from donors who have provided written informed consent. When using material from deceased organ donor those with legal authority were consented. Subject characteristics are presented in **Data file S1**.

### Healthy controls

A total of 234 healthy volunteers (56% females, 44% males, age range 1–85y) participated in the study as unpaid healthy controls. All denied having any chronic or acute disease.

### Temporal experiment cohort

15 healthy volunteers gave blood each week for 6 weeks (9 females, 6 males, age 21-68y).

### Vaccination cohort and determination of anti-hemagglutinin antibody titers

92 healthy volunteers that received the annual influenza vaccination (55 females, 37 males, age range 20-73y) gave blood samples a day before vaccination, and after 3, 7 and 28 days (± 2 days).

The anti-hemaggutinin antibody titers were determined using hemagglutination inhibition (HI) assay. Serum samples obtained from vaccinated and non-vaccinated individuals were stored at −20°C until tested treated with receptor destroying enzyme (RDE) (Sigma C8772, diluted 1:4), for 16 h prior to heat inactivation (30 min, 56°C). Absorption with erythrocytes was performed to remove non-specific hemagglutination, in accordance with a modified WHO protocol(Rowe et al., 1999). Serial two-fold dilutions (1:20–1:2560) of sera in 25 μl PBS were prepared in V-shaped well plates, and an equal volume of four hemagglutinin (HA) units of viral antigen was added. The mixture was then incubated at room temperature for 1 h. Fifty microliters of 0.5% chicken erythrocytes suspended in PBS, were added to the wells, and mixed by shaking the plates on a mechanical vibrator. Agglutination patterns were read after 30 min and the hemagglutination inhibition (HI) titer was defined as the reciprocal of the last dilution of serum that fully inhibited hemagglutination. The cut-off value selected for a positive result was 1:40. The influenza antigens for 2018-19 and 2019-20 winter seasons were supplied by the WHO.

### EoE cohort

21 active EoE patients, 24 EoE patients in remission and 14 controls where recruited to the study at Cincinnati Children’s Hospital. Diagnosis of EoE patients was made based on an histological biopsy taken from the distal esophageal tissue.

### Lymphoma cohort

27 newly diagnosed lymphoma patients that came for treatment in the hematological daycare unit in Hadassah medical center were recruited to the study in two cohorts (17 patients in cohort #1, 10 patients in cohort #2). Diagnosis was made by PET-CT.

### Sample collection and processing

Blood samples were collected by routine veinipuncture in 10 mL EDTA Vacutainer® tubes or Streck® blood collection tubes and stored at room temperature for up to 4 hours or 5 days, respectively. Tubes were centrifuged at 1,500×g for 10 minutes at 4°C (EDTA tubes) or at room temperature (Streck tubes). The supernatant was transferred to a fresh 15 mL conical tube without disturbing the cellular layer and centrifuged again for 10 minutes at 3000×g. The supernatant was collected and stored at -80°C.

cfDNA was extracted from 2-4 mL of plasma using the QIAsymphony liquid handling robot (Qiagen). cfDNA concentration was determined using Qubit double-strand molecular probes kit (Invitrogen) according to the manufacturer’s instructions.

DNA derived from all samples was treated with bisulfite using EZ DNA Methylation-Gold™ (Zymo Research), according to the manufacturer’s instructions, and eluted in 24μl elution buffer.

### Immune cell and tissue isolation and processing

PBMCs from a healthy individual where isolated using ficoll-paque density gradient (Miltenyi Biotec). CD4+ T-cells, CD8+ T-cells, CD19+ B-cells and Nk CD56+ cells were positively selected using magnetic MicroBeads. Monocytes where negatively selected (Miltenyi Biotec) as instructed by the manufacturer. Regulatory T-cells (CD4+,CD25+,FOXP3+, 28.5% purity) where purchased from Astarte biologics. Neutrophils and eosinophils where isolated based on a previously published protocol(Hartman et al., 2001; Sagiv et al., 2016) Genomic DNA from other tissues was purchased as previously described(Lehmann-Werman et al., 2016; Zemmour et al., 2018).

### Selection of immune cell methylation markers

immune-cell-specific methylation candidate biomarkers were selected using comparative methylome analysis, based on publicly available datasets(Moss et al., 2018a), to identify loci having more than five CpG sites within 150bp, with an average methylation value for a specific cytosine (present on Illumina 450K arrays) of less than 0.3 in the specific immune cell type and greater than 0.8 in over 90% of tissues and other immune cells. From our previously-described atlas of human tissue-specific methylomes(Lehmann-Werman et al., 2016), we identified ∼50 CpG sites that are unmethylated in specific immune-cells and methylated in all other major immune cells and tissues. We selected two to three of these sites for Neutrophils (i.e., NEUT1, NEUT2, NEUT3), Monocytes (i.e., MONO1, MONO2), Eosinophils (i.e., EOSI1, EOSI2, EOSI3), B-cells (i.e., B-CELL1, B-CELL2) T-cells (i.e., T-CELL1, T-CELL2), CD8 T-cells (CD8A, CD8B), Regulatory T-cells (TREG1, TREG2) and designed primers to amplify ∼100bp fragments surrounding them using the multiplex two-step PCR amplification method(Neiman et al., 2020). Marker coordinates and primer sequences are provided in **Supplementary Table S1.**

### PCR

To efficiently amplify and sequence multiple targets from bisulfite-treated cfDNA, we developed a two-step multiplexed PCR protocol. In the first step, up to 30 pairs primer pairs were used in one PCR reaction to amplify regions of interest from bisulfite-treated DNA, independent of methylation status. Primers were 18-30 base pairs (bp) with primer melting temperature ranging from 58-62°C. To maximize amplification efficiency and minimize primer interference, the primers were designed with additional 25bp adaptors comprising Illumina TruSeq Universal Adaptors without index tags. All primers were mixed in the same reaction tube. For each sample, the PCR reaction was prepared using the QIAGEN Multiplex PCR Kit according to manufacturer instructions with 7μl of bisulfite treated cfDNA. Reaction conditions for the first round of PCR were: 95°C for 15 minutes, followed by 30 cycles of 95°C for 30 seconds, 57°C for 3 minutes and 72°C for 1.5 minutes, followed by 10 minutes at 68°C.

In the second PCR step, the products of the first PCR reaction were treated with Exonuclease I (ThermoScientific) for primer removal according to the manufacturer instructions. <u>C</u>leaned PCR products were amplified using one unique TruSeq Universal Adaptor primer pair per sample to add a unique index barcode to enable sample pooling for multiplex Illumina sequencing. The PCR reaction was prepared using 2x PCRBIO HS Taq Mix Red Kit (PCR Biosystems) according to manufacturer instructions. Reaction conditions for the second round of PCR were: 95°C for 2 minutes, followed by 15 cycles of 95°C for 30 seconds, 59°C for 1.5 minutes, 72°C for 30 seconds, followed by 10 minutes at 72°C. The PCR products were then pooled, run on 3% agarose gels with ethidium bromide staining, and extracted by Zymo GEL Recovery kit.

### Next Generation Sequencing

Pooled PCR products were subjected to multiplex next-generation sequencing (NGS) using the MiSeq Reagent Kit v2 (Illumina) or the *NextSeq* 500/550 v2 Reagent Kit (Illumina). Sequenced reads were separated by barcode, aligned to the target sequence, and analyzed using custom scripts written and implemented in R. Reads were quality filtered based on Illumina quality scores. Reads were identified as having at least 80% similarity to the target sequences and containing all the expected CpGs. CpGs were considered methylated if “CG” was read and unmethylated if “TG” was read. Proper bisulfite conversion was assessed by analyzing methylation of non-CpG cytosines. We then determined the fraction of molecules in which all CpG sites were unmethylated. The fraction obtained was multiplied by the concentration of cfDNA measured in each sample, to obtain the concentration of tissue-specific cfDNA from each donor.

### Deconvolution

WGBS data were converted to an array-like format by calculating the average methylation at 7,890 CpGs from the Moss et al methylation atlas(Moss et al., 2018a). We then ran the deconvolution algorithm (https://github.com/nloyfer/meth_atlas) for each WBC and cfDNA sample.

### Statistics

To assess the correlation between groups we used Pearson’s correlation test. To determine the significance of differences between groups we used a non-parametric two-tailed Mann–Whitney test. For multiple comparisons, a Kruskal-Wallis multiple comparison test was used. P-value was considered significant when <0.05. To detect outliers in the healthy population we applied a multiple outlier detection ROUT-test (Q=5%)(Motulsky and Brown, 2006). Samples that were detected as outliers where excluded. All Statistical analyses were performed with GraphPad Prism 8.4.3.

### Intra-individual and inter-individual variation

Intra-individual coefficient of variation for each immune cell type in CBC, whole blood and cfDNA was calculated for each person across six different time points. The Inter-individual coefficient of variation for each immune-cell type was calculated for each time point across all individuals. The average of the intra-individual coefficient of variation was calculated. To prevent a bias due to difference in sample size (intra-individual variation, 6 time points; inter-individual variation, 15 individuals), we used R (version3.6.1) to sample all different combinations of a randomly selected 6 person group and calculated the inter-individual coefficient of variation. Coefficients of variation of the different combinations were averaged.

### Author contributions

Conceptualization-I.F-F, T.K, B.G, R.S, Y.D; Investigation-I.F-F, S.P, B-L.O, A.K, Ju.M, A.P, N.L, Jo.M, D.C, Y.D, N.F, J.M.C, M.R, A.J, G.C; Resources- M.M, M.E.R, D.L, T.K; Writing – I.F-F, B.G, R.S, Y.D.

## Acknowledgements

We thank Daniela Beller from the Maccabi TIPA biobank for providing data on blood cell counts, and the many volunteers who donated blood for this study. We thank Shai Shen-Orr for critical reading of the manuscript, and Nir Friedman, Ron Milo, Ron Sender and Mordechai Slae for fruitful discussions. We thank Idit Shiff and Abed Nasseredin from the Genomics lab of the Core Research Facility (CRF) at The Faculty of Medicine, The Hebrew University of Jerusalem. For their support in DNA and sequencing analysis.

This work was supported by grants from Human Islet Research Network (HIRN UC4DK116274 and UC4DK104216 to R.S and Y.D); JDRF (3-SRA-2014-38 and 1-SRA-2019-705), Ernest and Bonnie Beutler Research Program of Excellence in Genomic Medicine, The Alex U Soyka pancreatic cancer fund, The Israel Science Foundation, the Waldholtz / Pakula family, the Robert M. and Marilyn Sternberg Family Charitable Foundation, the Helmsley Charitable Trust, Grail and the DON Foundation (to Y.D). Y.D holds the Walter and Greta Stiel Chair and Research grant in Heart studies. I.F-F received a fellowship from the Glassman Hebrew University Diabetes Center.

---

**Supplementary Table S1.**
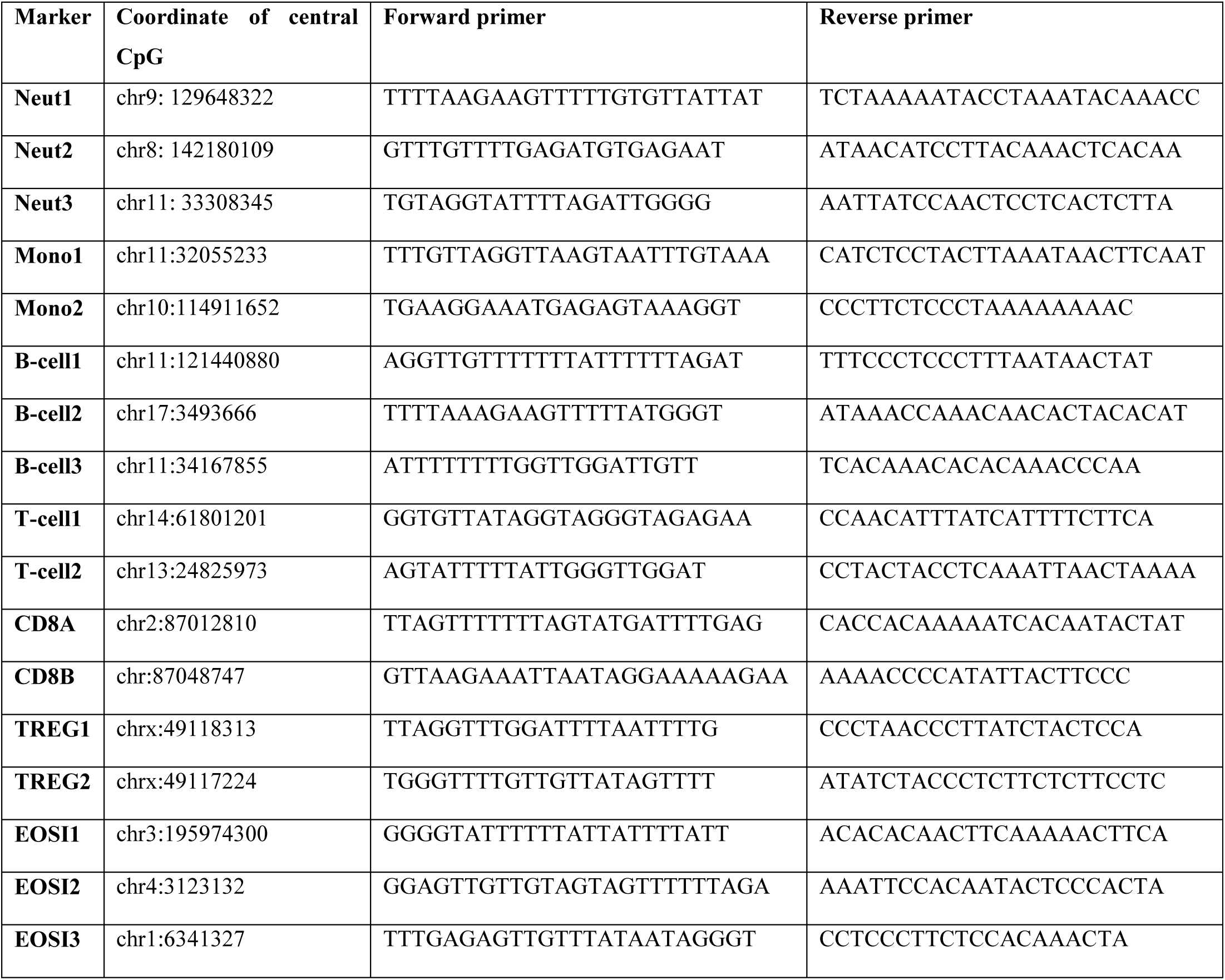
Genomic coordinates of immune cell type-specific methylation markers used in this study, and primer sequences used to amplify these loci after bisulfite conversion.

**Data file S1: Blood donor characteristics** (see excel file).

**Supplemental Figure S1:**
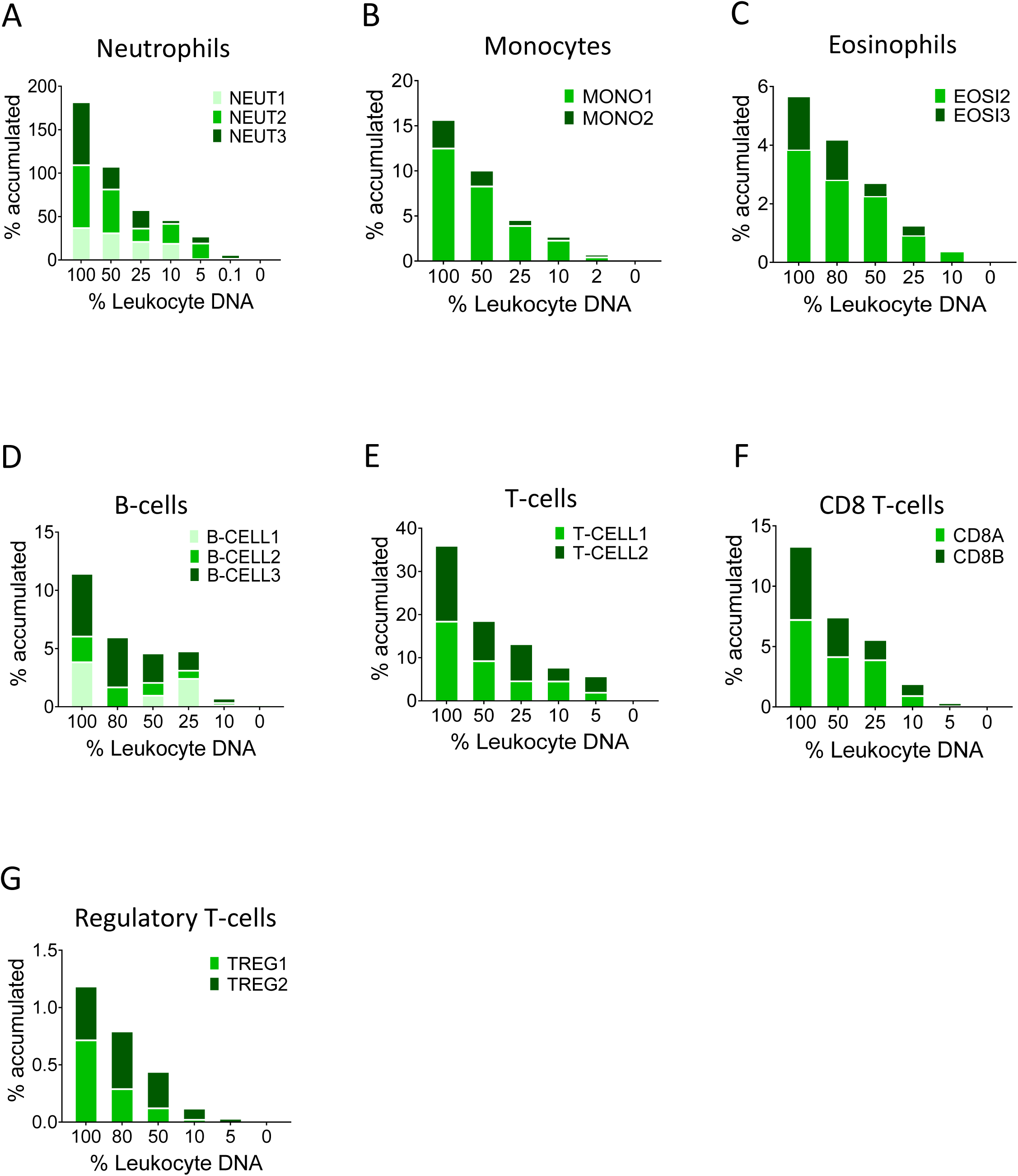
Identification of specific immune-cell DNA methylation markers. **A-G**, Spike in experiments for the assessment assay sensitivity. Human leukocyte DNA was mixed with DNA from HEK293 cells in the indicated proportion. The fraction of DNA from each immune type cell was assessed using the indicated methylation markers. The Y axis shows cumulative values for percentage contribution from each marker, to underscore the signal from each individual marker.

**Supplemental Figure S2:**
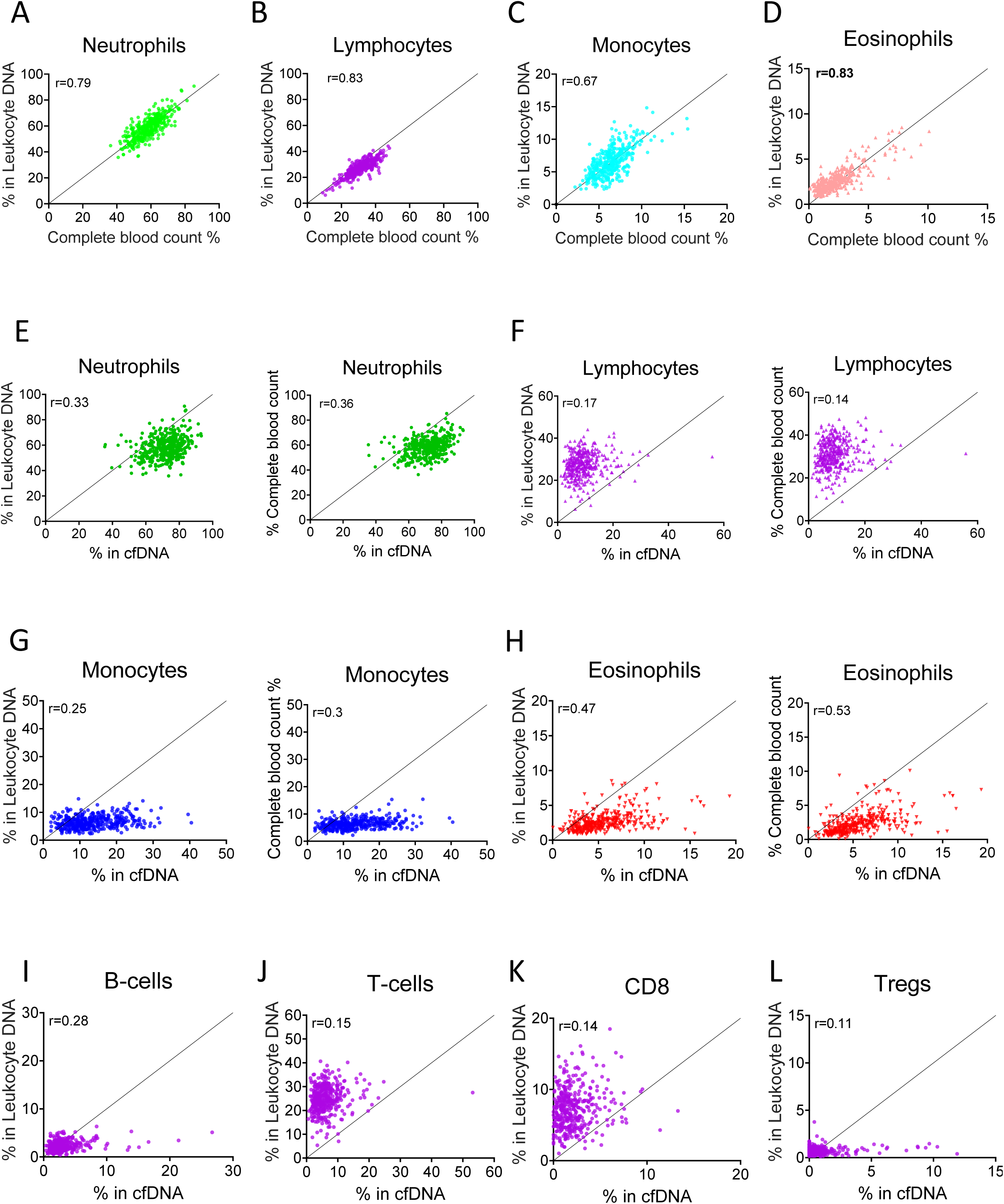
Levels of immune cell-derived cfDNA are not correlated with the counts of these cells in circulation, and reflect cell type-specific turnover rates. Plasma and blood samples (n=392) were obtained from 79 healthy individuals that vary in age and gender. Methylation-based immune markers were determined on genomic DNA from whole blood, and on plasma cfDNA. Standard CBC values were also obtained. **A-D,** correlation between CBC and methylation markers in cfDNA. **E-H**, correlation between cfDNA methylation signals and whole blood methylation signals (left graph in each panel) or CBC (right graph in each panel). **I-L**, correlation between cfDNA methylation signals whole blood methylation signals for cell types that are not scored in CBC.

**Supplemental Figure S3:**
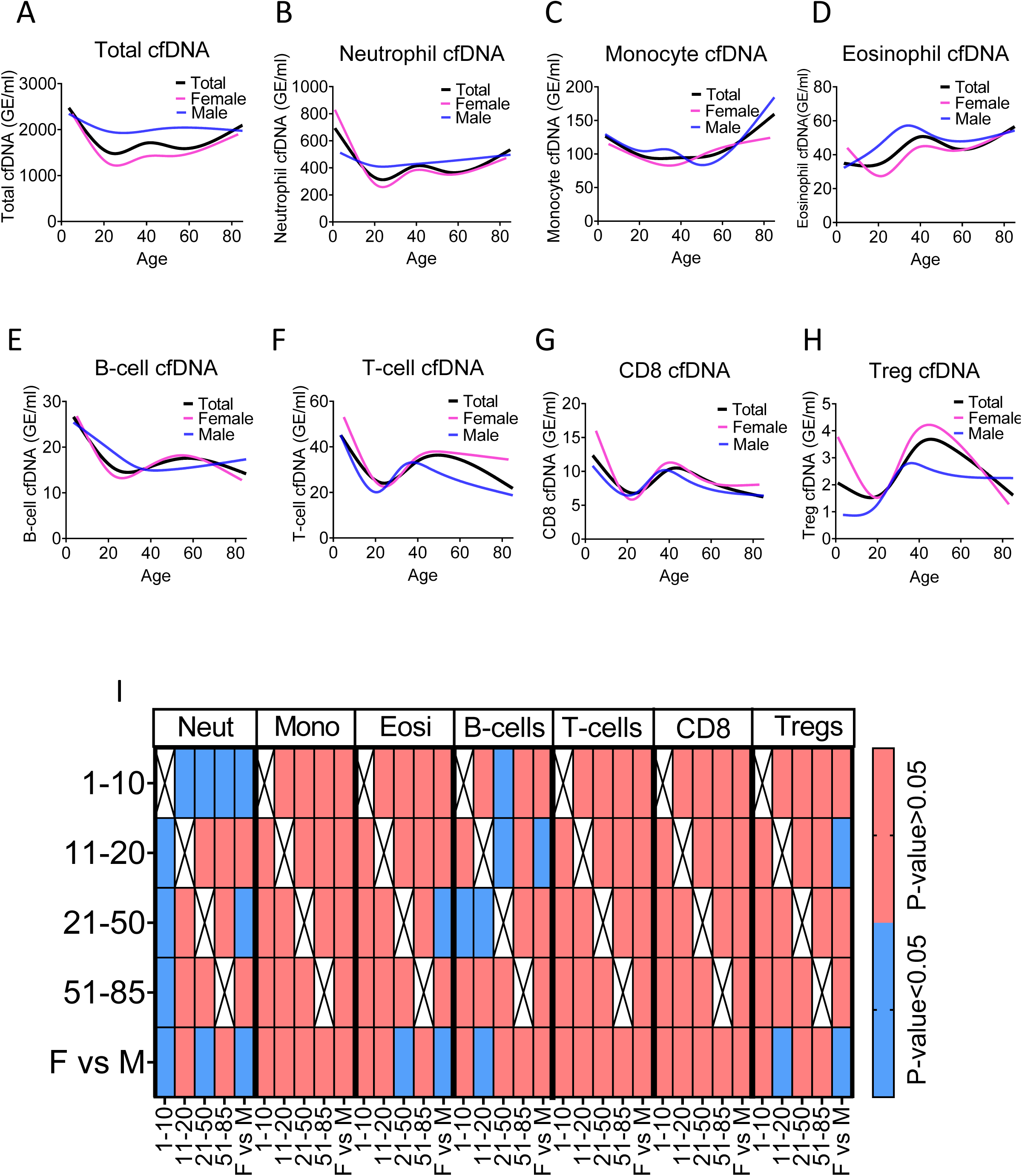
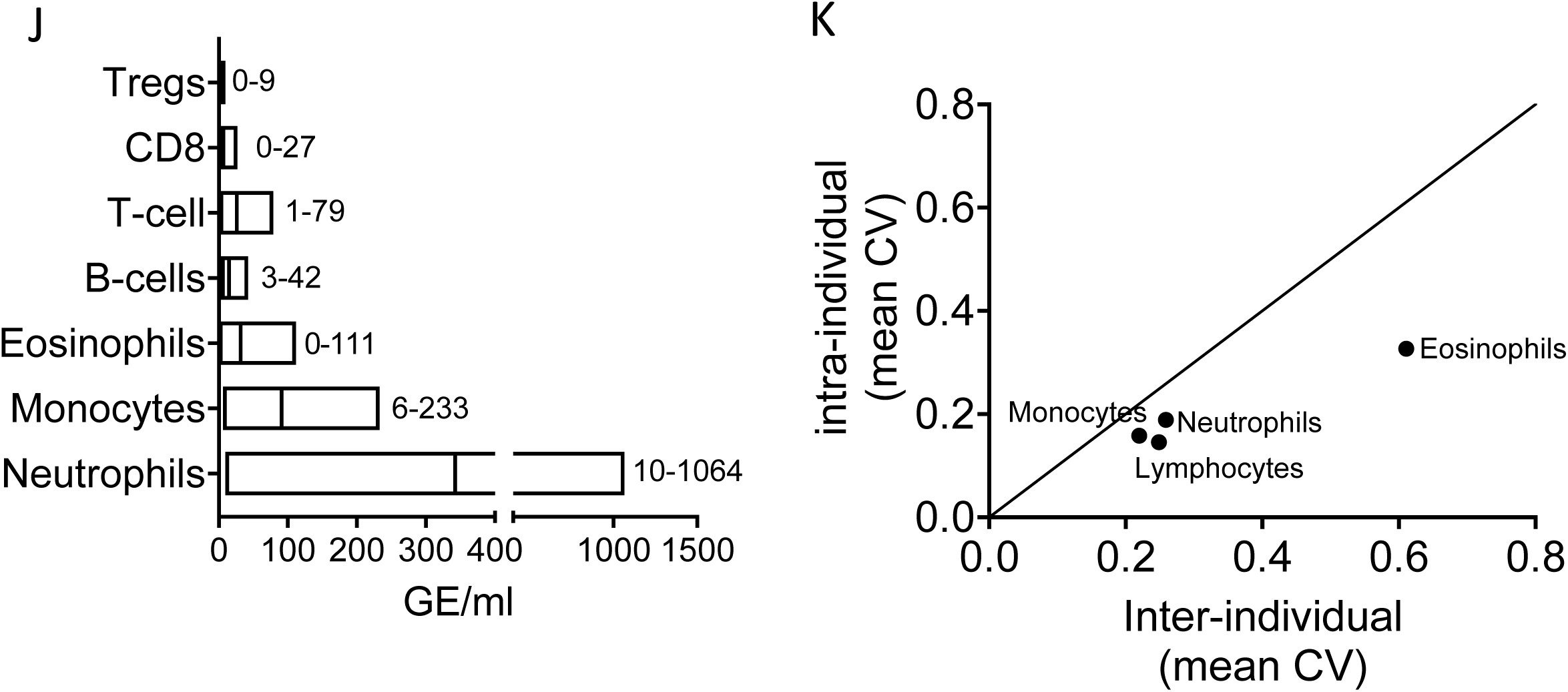
Immune derived cfDNA characterization in a healthy population. **A-H**, total and immune derived cfDNA was characterized in healthy individuals (n=227) as a spline curve fit showing the trend of the data across different age groups (black line), with a breakdown to gender (female pink lines, male blue lines) for the indicated immune cell types. **I,** a categorical heat-map showing significant (blue, p-value<0.05) and non-significant (red, p-value>0.05, Kruskal-wallis) differences in cfDNA values for each immune cell type across different age groups and between genders. Age groups were 1-10 years (n=21), 11-20 (n=30), 21-50 (n=134), and >50 (n=37). **J,** the normal cfDNA range for each cell type: neutrophils (mean=390, range 10-1064 GE/ml), monocytes (mean=101, range 6-233 GE/ml), eosinophils (Mean=38, range 0-111 GE/ml) T cells (Mean=30, range1-79) B-cells (mean=17, range 3-42 GE/ml) CD8 T-cells, (mean=8, range 0-27 GE/ml) and Tregs (mean=2, range 0-9 GE/ml) based on samples from 227 individuals excluding outliers using the ROUT outlier test (Q=5%). **K,** XY scatter plot showing the average of inter-individual coefficient of variation (X-axis) and intra-individual coefficient of variation (Y-axis) for the fraction of cells from each immune cell types based on CBC. Black line represents perfect correlation between inter- and intra-individual variation. Dots below the black line represent greater inter-individual variation and dots above represent greater intra-individual variance.

**Supplemental Figure S4:**
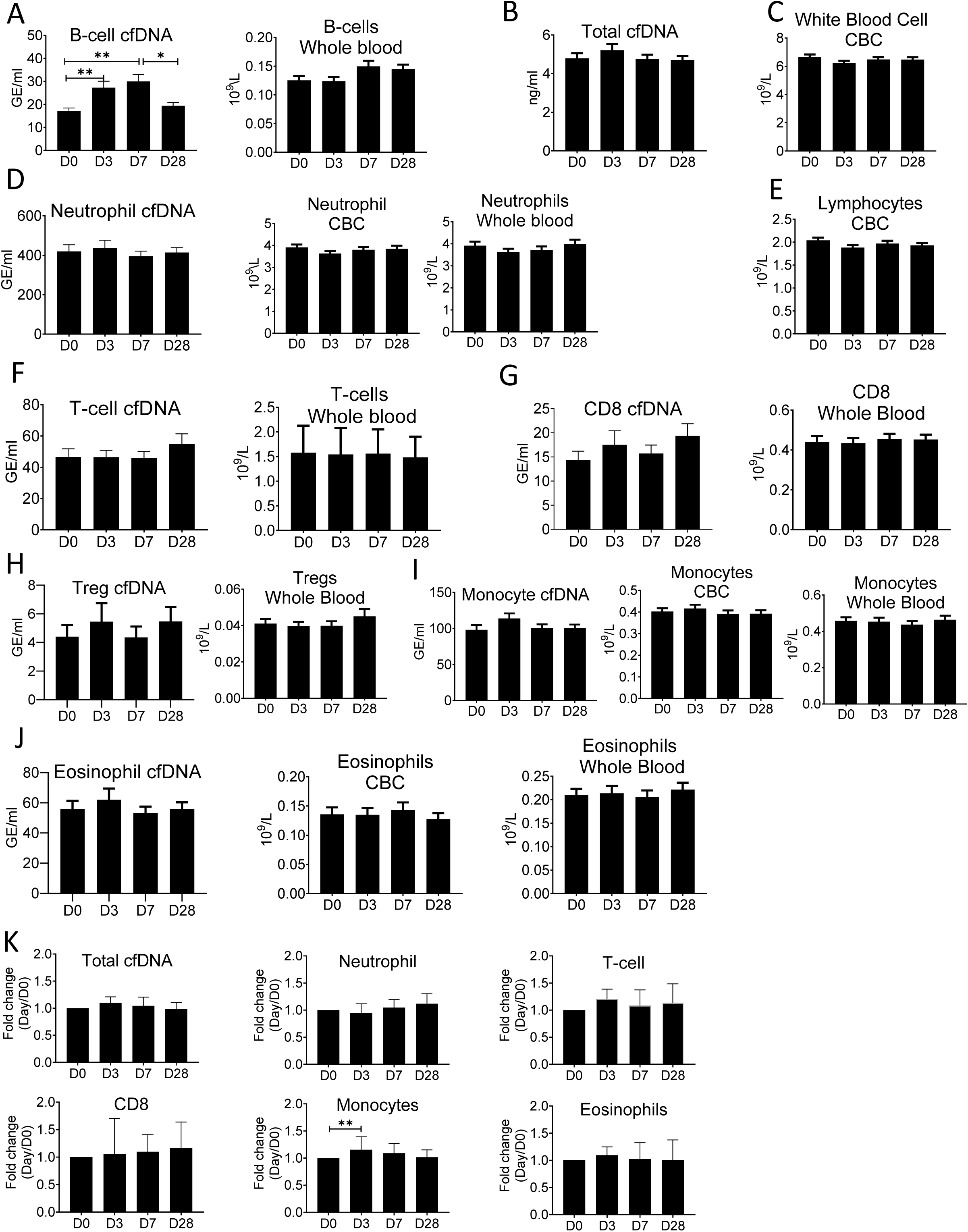
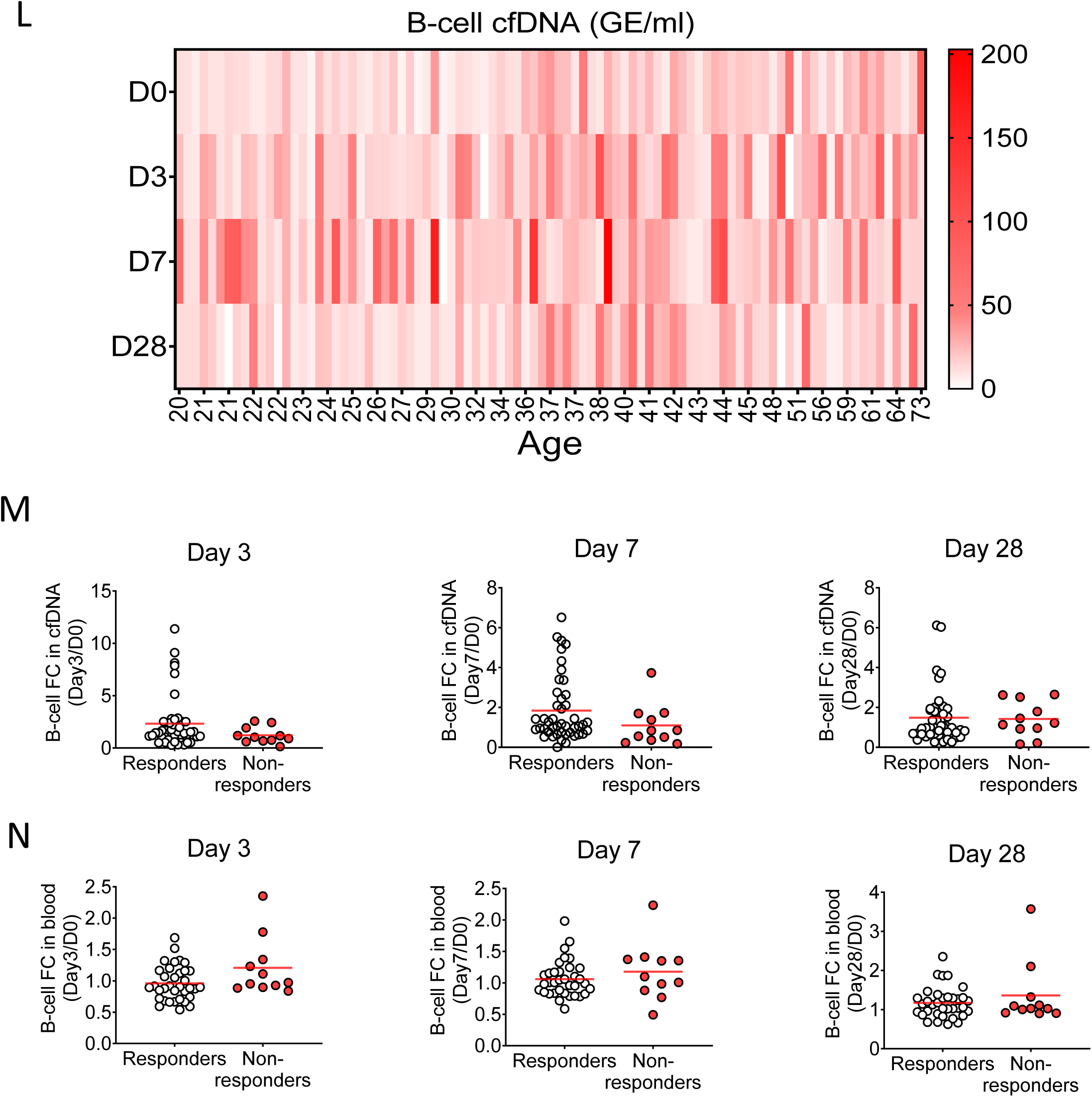
Specific elevation of B-cell derived cfDNA after influenza vaccination, prior to changes in cell counts and in correlation to efficacy of response to vaccination. Plasma and blood samples were taken from volunteers (n=92) receiving the annual influenza vaccination. Samples were obtained on the day before vaccination (D0), and at days 3,7 & 28 (D3, D7, D28). Immune methylation markers where tested on both cfDNA and whole blood. Statistical significance is calculated relative to the levels at D0. **A,** B-cell derived cfDNA markers are elevated on day 3 and day 7 (p-value=0.0002) after influenza vaccination. Circulating B-cell counts did not change significantly at the population level (p-value=0.078). **B,** Total cfDNA (p-value=0.9). **C,** White blood cells count (p-value=0.28). **D,** Neutrophil DNA in plasma (p-value=0.65) and whole blood (p-value=0.53), and neutrophil counts in CBC (p-value=0.47). **E,** Lymphocyte counts in CBC (p-value=0.26). **F,** T-cell DNA in plasma (p-value=0.58) and whole blood (p-value=0.84). **G,** CD8 T-cell DNA in plasma (p-value=0.5) and whole blood (p-value=0.83). **H**, Treg DNA in plasma (p-value=0.78) and whole blood (p-value=0.85). **I,** Monocyte DNA in plasma (p-value=0.22) and whole blood (p-value=0.68), and monocyte counts in CBC (p-value=0.79). **J,** Eosinophil DNA in plasma (p-value=0.76) and whole blood (p-value=0.81), and eosinophil counts in CBC (p-value=0.93). **K**, cfDNA measurements normalized to baseline at D0 of each individual: total cfDNA (p-value=0.37), neutrophil cfDNA (p-value=0.57), T-cell cfDNA (p-value=0.69), CD8 cfDNA (p-value=0.48), monocyte cfDNA (p-value=0.01), and eosinophil cfDNA (p-value=0.88).(Kruskal-wallis). **L,** a heat map showing the level of B-cell cfDNA in each individual following vaccination. **M-N,** normalized B-cell derived cfDNA **(M)** and normalized circulating B-cells (**N**) in responders versus non-responders on days 3, 7 and 28. There are no statistically significant differences between the groups (Mann-Whitney test). Note that when the peak value of B cell cfDNA in blood is considered (at any day after vaccination), responders have a stronger signal, as shown in main Figure 4.

**Supplemental Figure S5:**
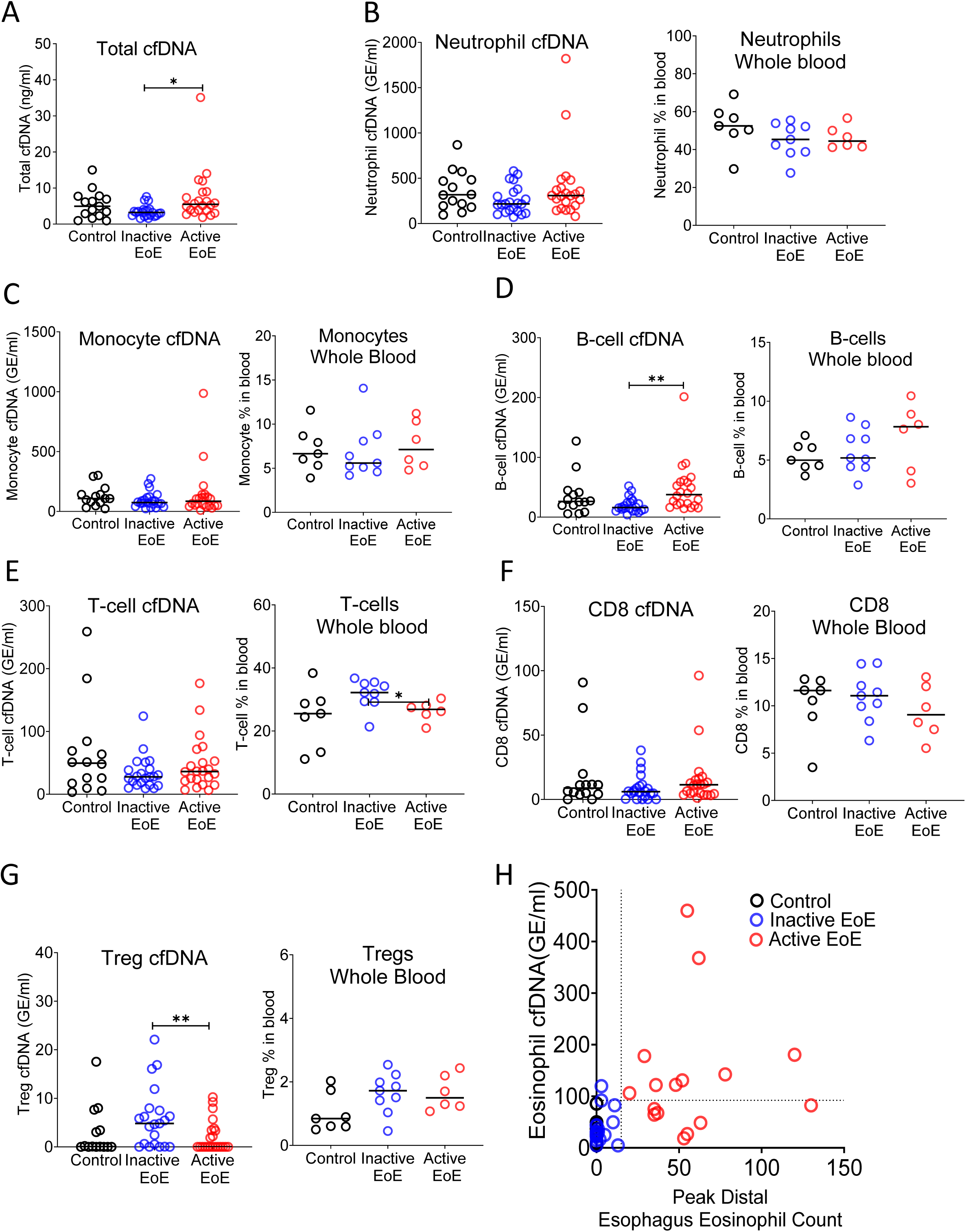
Selective elevation of eosinophil-derived cfDNA in patients with Eosinophilic Esophagitis. Plasma was collected from three groups: healthy controls (n=14), inactive EoE (n=24), and active EoE (n=21). Some of the patients had also PBMCs collected: healthy controls (n=7), inactive EoE (n=9), and active EoE (n=6). **A,** Total cfDNA levels were significantly higher in active EoE compared to inactive EoE (p-value=0.01). **B,** neutrophil derived cfDNA (p-value=0.19) and neutrophil DNA in whole blood (p-value=0.23) are not different in EoE. **C,** monocyte cfDNA (p-value=0.45) and monocytes DNA in whole blood are not different in EoE (p-value=0.75). **D,** B-cell derived cfDNA is significantly higher in active EoE compared with inactive EoE (p-value=0.0021), but B-cell DNA in whole blood is not significantly different in EoE (p-value=0.47). **E,** T-cell derived cfDNA is not significantly different between the groups (p-value=0.48). However the fraction of T-cell DNA in whole blood is significantly higher in inactive EoE compared with active EoE (p-value=0.0454). **F,** there is no significant difference in CD8 derived cfDNA (p-value=0.4) and in CD8-derived DNA in whole blood among EoE patients (p-value=0.6). **G,** Treg cfDNA levels are significantly higher in inactive EoE compared with active EoE (p-value=0.009) while Treg DNA in whole blood is not different between patients and controls (p-value=0.15, Kruskal-Wallis test). **H,** XY Scatter plot for Eosinophil derived cfDNA levels versus Peak distal Esophagus Eosinophil count, in healthy controls and in patients with inactive or active EoE. Dashed lines indicate thresholds for negative and positive Peak distal Esophagus Eosinophil count (HPF), and negative and positive Eosinophil cfDNA.

**Supplemental Figure S6:**
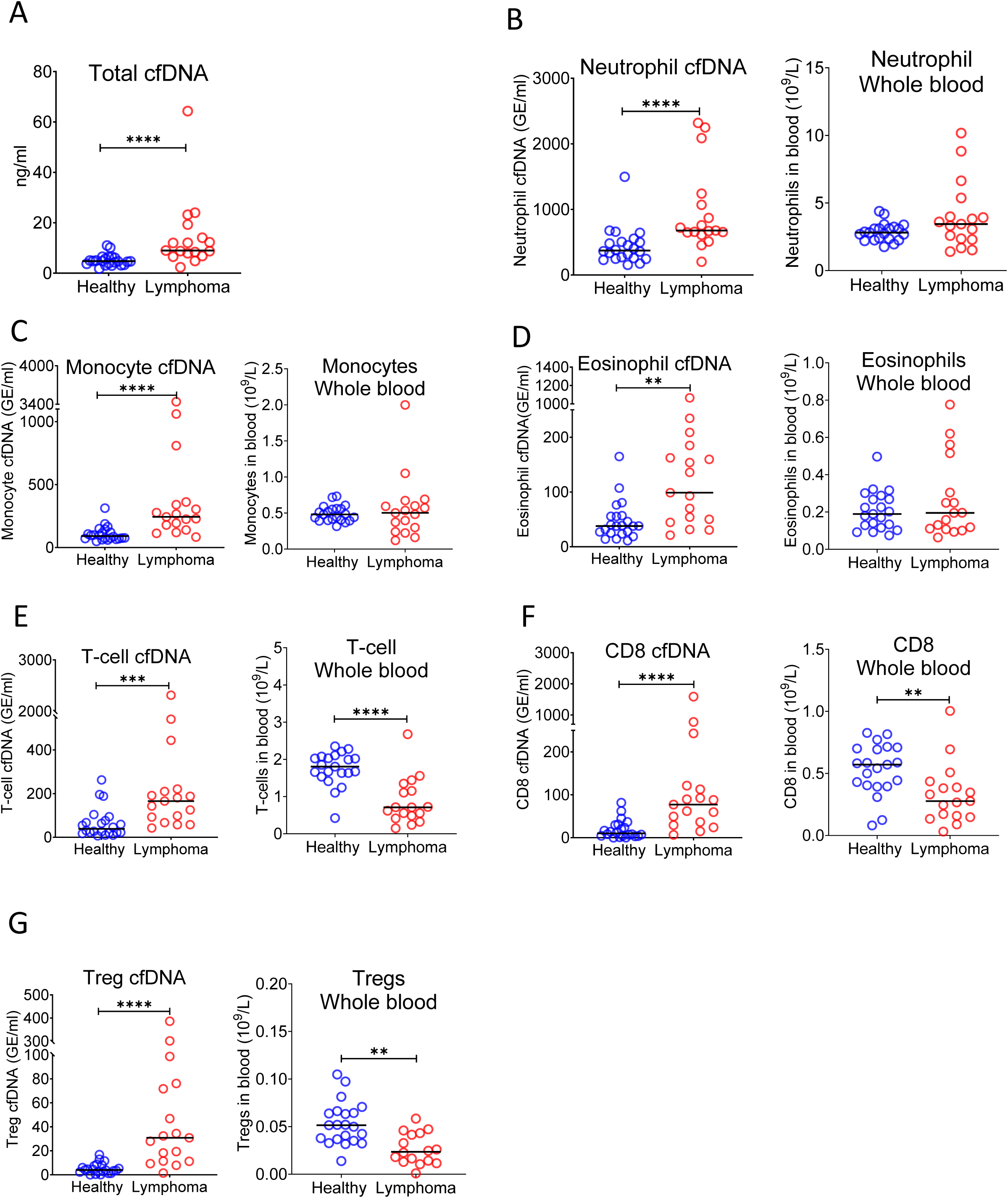
Immune derived cfDNA in lymphoma patients. Plasma and blood were collected from healthy controls (n=22) and patients with lymphoma (n=17). **A,** Total cfDNA levels are significantly higher in lymphoma patients compared with healthy controls (p-value<0.0001 Mann–Whitney test). **B**, neutrophil-derived cfDNA levels are higher in lymphoma patients compared with controls (p-value<0.0001), but neutrophil DNA levels in whole blood are not different between patients and controls (p-value=0.17). **C,** monocyte-derived cfDNA levels are elevated in lymphoma patients (p-value<0.0001), but monocyte DNA levels in whole blood are not different between patients and controls (p-value=0.95). **D,** eosinophil-derived cfDNA are elevated in lymphoma patients (p-value<0.0019) but eosinophil DNA in whole blood is not elevated (p-value=0.86). **E,** T-cell derived cfDNA levels are elevated in lymphoma (p-value=0.003) while T-cell DNA is reduced in whole blood (p-value<0.0001). **F**, CD8 T-cell derived cfDNA levels are elevated in lymphoma (p-value<0.0001) while CD8 T-cell DNA is reduced in whole blood (p-value=0.003). **G,** Treg-derived cfDNA levels are elevated in lymphoma patients (p-value<0.0001) while Treg DNA in whole blood is reduced (p-value=0.0026, Mann– Whitney test).

**Supplemental Figure S7:**
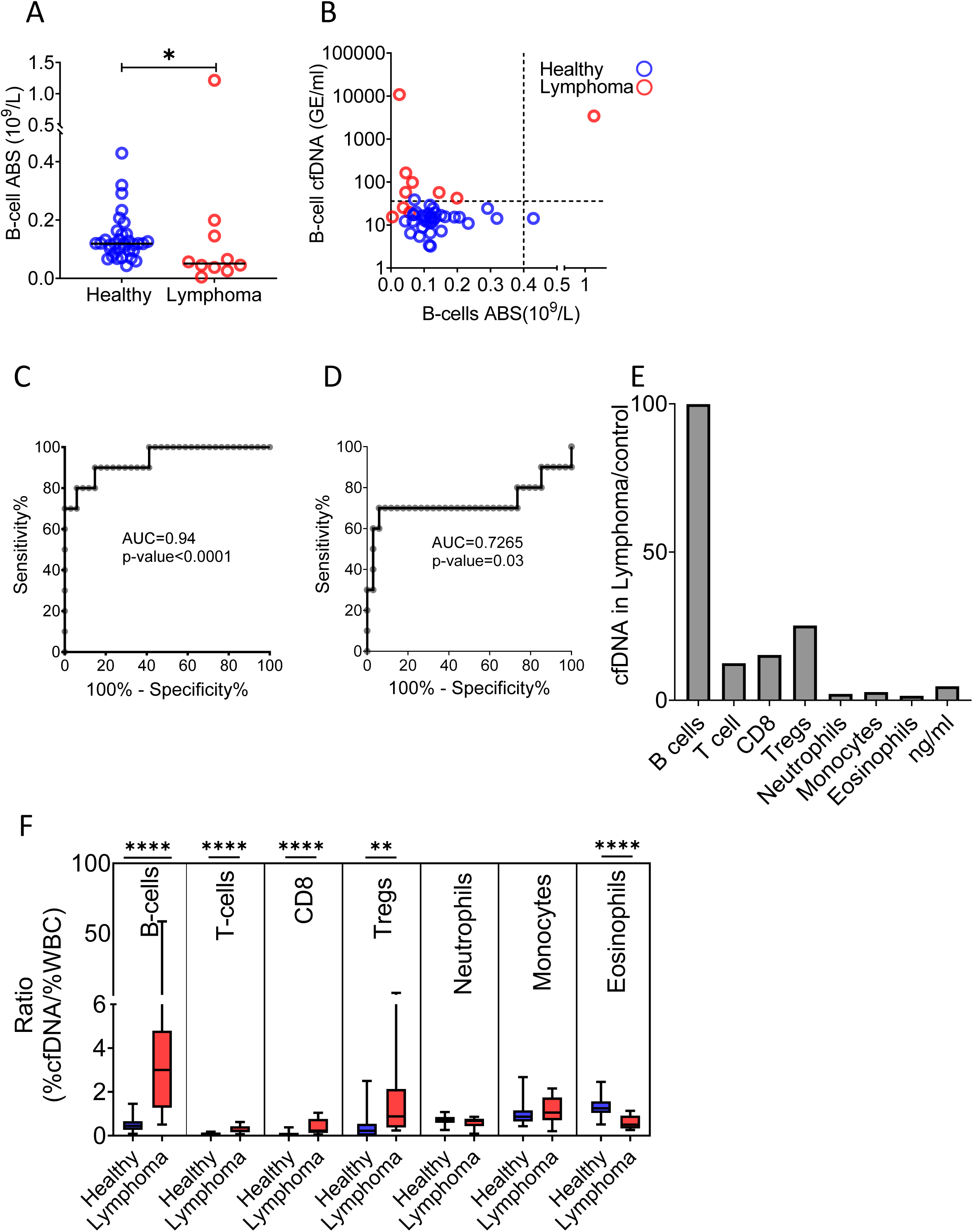
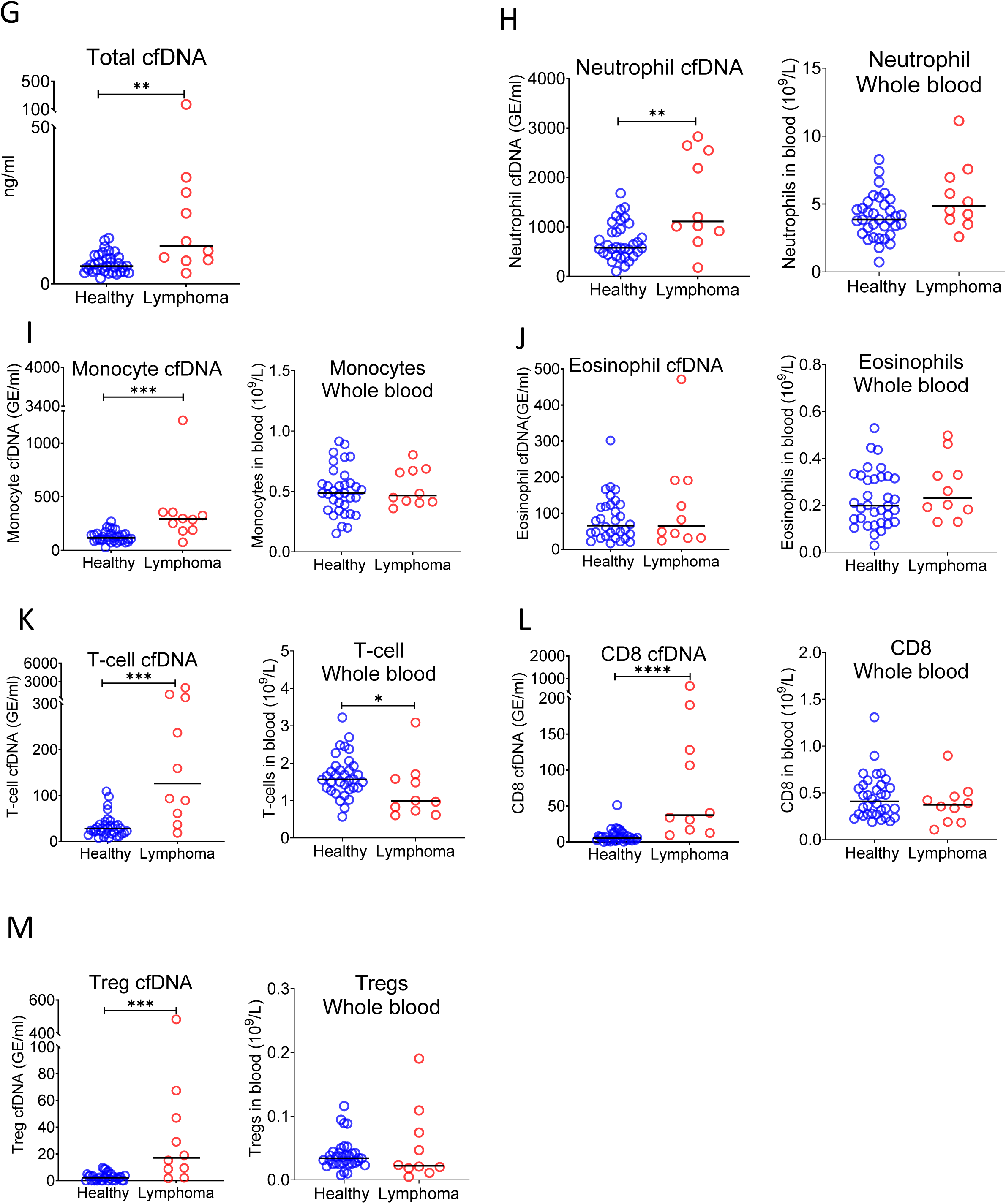
cfDNA analysis of a second cohort of lymphoma patients. Blood was collected from a 2nd cohort of donors including 34 healthy controls and 10 patients with lymphoma. **A,** B-cell counts (p-value=0.03, Mann-Whitney). **B,** XY Scatter plot for B-cell derived cfDNA levels versus B-cell absolute counts. Dashed lines indicate healthy baseline levels of B-cell absolute counts and B-cell cfDNA. **C,** ROC curve for the diagnosis of lymphoma based on B-cell cfDNA levels in healthy subjects and patients with B cell lymphoma. **D,** ROC curve for diagnosis of lymphoma based on B-cell counts. **E,** immune cell type-specific cfDNA in lymphoma patients and healthy controls (mean lymphoma/ mean control). **F,** the ratio between the percentage of cfDNA from a given immune cell type and the percentage of cells from this population in blood according to CBC, in each donor of the 2nd cohort (healthy blue bars; lymphoma red bars). Boxes represent 25th and 75th percentiles around the median, whiskers span min to max. **G,** Total cfDNA levels are significantly higher in lymphoma patients compared with healthy controls (p-value=0.0027 Mann–Whitney test). **H,** neutrophil-derived cfDNA levels are higher in lymphoma patients compared with controls (p-value=0.0067), but neutrophil DNA levels in whole blood are not different between patients and controls (p-value=0.07). **I,** monocyte-derived cfDNA levels are elevated in lymphoma patients (p-value=0.001), but monocyte DNA levels in whole blood are not different between patients and controls (p-value=0.71). **J,** eosinophil-derived cfDNA and whole blood are not significantly higher in lymphoma patient’s vs healthy controls (p-value=0.63; p-value=0.28). **K,** T-cell derived cfDNA levels are elevated in lymphoma (p-value=0.0001) while T-cell DNA is reduced in whole blood (p-value=0.036). **L,** CD8 T-cell derived cfDNA levels are elevated in lymphoma (p-value<0.0001) while CD8 T-cell DNA in whole blood is not different between patients and controls (p-value=0.37). **M,** Treg-derived cfDNA levels are elevated in lymphoma patients (p-value=0.0001) while Treg DNA is not significantly different in whole blood (p-value=0.41, Mann–Whitney test).

## Notes

### Competing Interest Statement

I.F-F., Ju.M., Jo.M, T.K, B.G., R.S. and Y.D. have filed patents on cfDNA analysis technology. A.J. and G.C. are employees of GRAIL. Other co-authors declare that no conflict of interest exists.

